# Structural basis for carbapenam *C*-alkylation by TokK, a B_12_-dependent radical SAM enzyme

**DOI:** 10.1101/2021.07.10.451909

**Authors:** Hayley L. Knox, Erica K. Sinner, Craig A. Townsend, Amie K. Boal, Squire J. Booker

## Abstract

Cobalamin- or B_12_-dependent radical *S*-adenosylmethionine (SAM) enzymes acting during carbapenem antibiotic biosynthesis carry out radical-mediated methyl transfers that underlie the therapeutic usefulness of these essential medicines. Here we present x-ray crystal structures of TokK, which are representative of this functional class, containing its two metallocofactors and determined in the presence and absence of carbapenam substrate. The structures give the first visualization of a cobalamin-dependent radical SAM methylase that employs the radical mechanism shared by a vast majority of these enzymes. The structures provide insight into the stereochemistry of initial C6 methylation and suggests that substrate positioning governs the rate of each methylation event.

**One Sentence Summary:** Structural insight into a cobalamin-dependent radical SAM methylase that performs three sequential radical-mediated methylations to install the C6 side chain of a carbapenem antibiotic.

---

Carbapenems are antibiotics of last resort in the clinic. Due to their potency and broad-spectrum activity, they are an increasingly important part of the arsenal to fight bacterial infections. Their continuing vital role is exemplified by Merck’s recent FDA approval for the use of the carbapenem imipenem in combination with cilastatin and relebactam to treat hospital-acquired and ventilator-associated bacterial pneumonia, a pressing problem during the ongoing COVID-19 pandemic (*1*). The C6 hydroxyethyl side chain distinguishes the clinically-used carbapenems from the other known classes of β-lactam antibiotics and is responsible for their low susceptibility to inactivation by β-lactamases owing to the occlusion of water from the β-lactamase active site (Figure 1A) (*2*).

**Figure 1.**
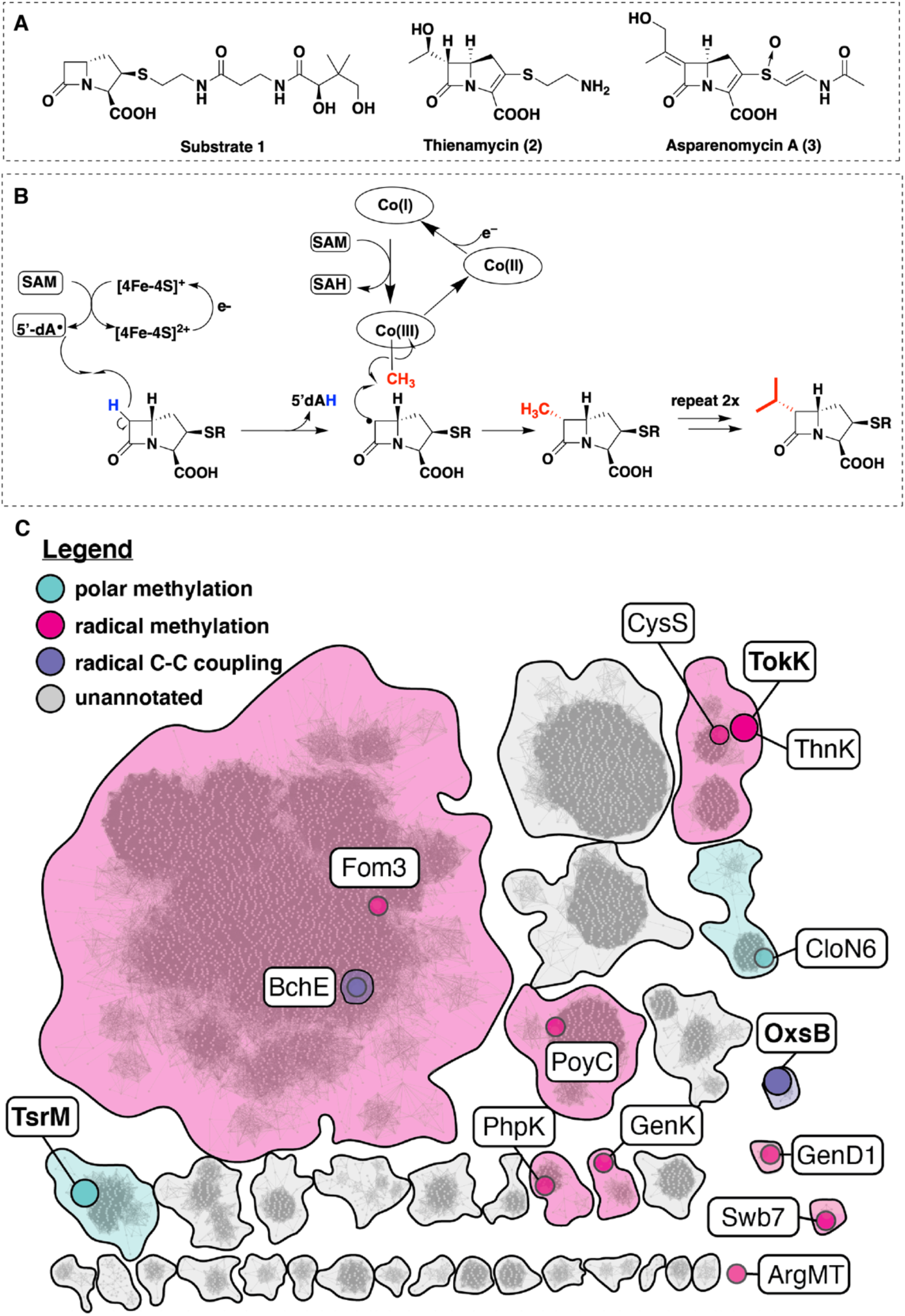
Cobalamin-dependent radical-mediated methylations in carbapenem biosynthesis. (**A**) Structure of (2*R*) pantetheinylated carbapenam precursor substrate **1** modified by ThnK and TokK. Related carbapenem natural products containing C6-alkyl substituents include thienamycin and asparenomycin A. (**B**) Proposed mechanism describing the three sequential methylations catalyzed by TokK. (**C**) An abbreviated sequence similarity network (SSN) of cobalamin-binding RS enzymes, highlighting selected sequence clusters and nodes. The full network (Figure S2) was generated from ∼11,000 annotated cobalamin-dependent RS enzymes with an alignment score of 65. Each node represents a single sequence or a set of sequences with >40% sequence identity. Sequence clusters and nodes are colored by predicted reaction mechanism (see legend in figure panel). Nodes containing functionally annotated sequences are indicated in color. Structurally characterized enzymes are represented by enlarged nodes and labeled in boldface.

During the biosynthesis of all known complex carbapenem natural products, the assembly of the C6 alkyl side chain is accomplished by a cobalamin (Cbl or B_12_)-dependent radical *S*-adenosylmethionine (SAM) enzyme (*3, 4*). These catalysts can perform serial methyl transfers with control of stereochemical outcome for each reaction. The Cbl-dependent radical SAM (RS) enzyme ThnK performs two sequential methyl transfers to its (2*R*)-pantetheinylated carbapenam substrate **1** during the biosynthesis of the paradigm carbapenem thienamycin (**2**) (*5*). An ortholog of ThnK, TokK from *Streptomyces tokunonensis* (ATCC 31569), constructs the C6 isopropyl chain of the carbapenem, asparenomycin A (**3**), by deploying three sequential methylations of **1 (**Figure 1B). This biosynthetic approach allows the producing organism to make a small “library” of alkylated analogues, which may deter the development of resistance in susceptible bacteria. This strategy may also be employed in the biosynthesis of cystobactamids, in which a similar Cbl-dependent radical SAM enzyme, CysS, performs successive methyl transfers (Figure S1) (*6*). Interestingly, despite low sequence identity (∼29%) between CysS and TokK or ThnK, all three proteins are located within the same node of a sequence similarity network (SSN) composed of approximately 11,000 cobalamin-binding RS enzymes (Figure 1C **and** S2). It is tempting to speculate that this colocalization might be driven by mechanistic similarities.

Carbapenem C6 alkyl chain construction requires stereoselective formation of carbon-carbon bonds between unactivated *sp*^3^*-*hybridized carbons. Cbl-dependent RS enzymes are the only known biological catalysts capable of such transformations. The Cbl-containing subfamily, depicted as an SSN in Figure 1C, is also one of the largest in the RS superfamily, a diverse group that functions in the biosynthesis of chlorophyll, novel lipids, and natural products with antiproliferative biological activity (*7–10*). While the majority of Cbl-dependent RS enzymes have unknown functions, those that have been characterized are generally, but not exclusively, methylases that act on carbon or phosphorus centers by using methylcobalamin (MeCbl) as an intermediate methyl donor. All RS enzymes, with a single known exception (*11–13*), reductively cleave SAM to generate methionine (Met) and a 5’-deoxyadenosyl 5’-radical (5’-dA•) (Figure 1B). The latter reactive intermediate typically initiates catalysis with a target substrate by abstracting a hydrogen atom. In B_12_-dependent RS methylases, the substrate radical attacks the methyl group of MeCbl, inducing homolytic cleavage of the cobalt-carbon bond to afford cob(II)alamin and the methylated product (Figure 1B). Upon dissociation of the methylated product, Met, and 5’-deoxyadenosine (5’-dAH), and rebinding of another molecule of SAM, cob(II)alamin is reduced to cob(I)alamin. Co(I), a supernucleophile, then acquires a methyl group from SAM to regenerate MeCbl (Figure 1B) (*9, 10*).

Two Cbl-dependent RS enzymes have been structurally characterized, TsrM and OxsB (Figure 1C **and** S2) (*11, 14*), involved in the biosynthesis of antibiotics thiostrepton A and oxetanocin A, respectively. Both enzymes are mechanistic outliers among Cbl-dependent RS enzymes, and are found in SSN nodes distinct from each other and from TokK (Figure 1C). OxsB uses cobalamin in an unknown fashion to catalyze a complex ring contraction of 2’-deoxyadenosine monophosphate (dAMP) (Figure S3) (*14*). TsrM methylates an *sp*^2^-hybridized carbon, C2, of L-tryptophan (Trp) by a polar mechanism (Figure S3) (*11*). TsrM is distinctive among all RS enzymes because it does not catalyze the formation of 5’-dA• during catalysis (*11*). Instead, TsrM uses SAM’s carboxylate moiety as an acceptor of the N1 proton of Trp during C2 electrophilic substitution by MeCbl (*11, 15*). Additionally, the structures of TsrM and OxsB have limitations that prevent full understanding of Cbl-dependent RS catalysis. The structure of OxsB lacks the dAMP substrate (*14*). TsrM has been co-crystallized with aza-SAM (a SAM analog) and Trp, but the Trp substrate is bound in an unproductive conformation, requiring computational docking to understand the structural basis for the reaction outcome. Herein we present x-ray structures of TokK, a Cbl-dependent RS enzyme that catalyzes canonical radical-dependent methylations, in both unliganded and substrate-bound forms. The substrate is bound in a catalytically competent conformation, providing the first view of functional substrate positioning in this important class of metalloenzymes and a structural rationale for the distribution of methylated products in TokK.

## RESULTS

TokK was crystallized under anoxic conditions in the presence of 5’-dAH and Met, the products of reductive SAM cleavage. Structures of this complex were solved in the absence and presence of substrate **1** to resolutions of 1.79 Å and 1.94 Å, respectively (Table S1). TokK shares in common with TsrM and OxsB an N-terminal Cbl-binding domain and a central RS domain containing a [4Fe-4S] cluster (Figure S4 **and** S5). A third C-terminal domain is distinct to TokK (Figure 2A **and** S6) (*11, 14*). While as-isolated TokK contains methylcobalamin (MeCbl), hydroxycobalamin (OHCbl), and adenosylcobalamin (AdoCbl), only OHCbl is observed bound to the N-terminal domain in the x-ray crystal structures. This assignment was confirmed by high-resolution mass spectrometry of dissolved crystals (Figure S7). In the structure of the TokK•OHCbl•5’-dAH•Met•substrate complex, which mimics the complex immediately prior to reaction with substrate, we observed clear *F*_o_-*F*_c_ electron density consistent with the shape and size of **1** in one of two monomers in the asymmetric unit (Figure 2B). In the second monomer, this electron density was also present, but it was of insufficient intensity for substrate modeling (Figure S8). In the chain with substrate bound, the pantetheine tail of **1** occupies a channel that leads from the surface of the protein into the active site (Figure 2A **and** S9). This cavity is formed at the interface of all three domains of TokK. The N-terminal Cbl-binding domain and unique C-terminal domain contribute the majority of interactions with the pantetheine unit (Figure 2A **and** S10). These include hydrophobic contacts, water-mediated H-bonding interactions, and direct polar contacts. For example, Asn515 in the C-terminal domain H-bonds to the terminal -OH of the pantetheine moiety, suggesting a key role for this domain in substrate recognition. In the N-terminal domain, the Cbl cofactor itself participates in a water-mediated contact to an amide carbonyl of **1**. This network involves one of the Cbl propionamide substituents and Tyr410 of the RS domain. The ß-lactam ring of **1** is buried deep within the RS domain and anchored by direct and water-mediated contacts to the C7 carbonyl and C3 carboxylate substituents. This mode of ß-lactam interaction resembles that of carbapenem synthase, the enzyme responsible for inversion of stereochemistry at C5 in simple carbapenems (Figure S11) (*16*). Both enzymes share use of H-atom abstraction chemistry to selectively target an unactivated C-H bond within the bicyclic ß-lactam core, consistent with their conserved substrate anchoring strategies. The C3 carboxylate of **1** H-bonds to Arg280 in the RS domain and Tyr652 in the C-terminal domain. These side chains move significantly from their positions in the structure without substrate bound (Figure 2A), and substitution of Arg280 to Gln results in near complete loss of activity (Figure 2C), confirming the importance of these side chains in substrate binding.

**Figure 2.**
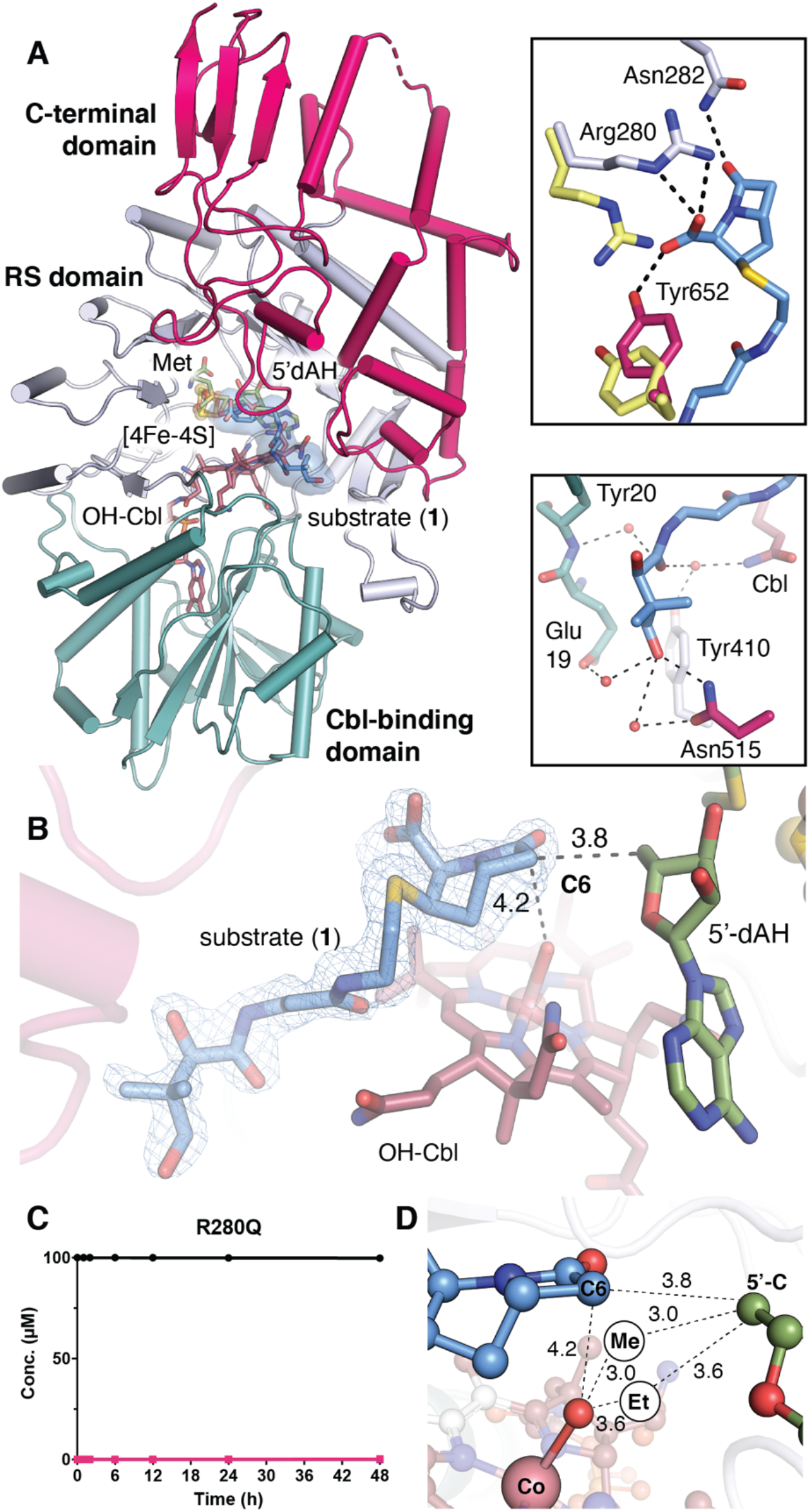
TokK binds its carbapenam substrate at the interface of three domains. **(A)** The overall structure of TokK (chain A) is illustrated as a ribbon diagram and colored by domain. The Cbl-binding Rossmann fold is shown in teal with the cobalamin cofactor in stick format and colored by atom type. The RS domain is shown in light blue. A [4Fe-4S] cluster is shown in orange and yellow spheres. 5’-dAH and Met coproducts are shown in stick format. The C-terminal domain is shown in pink. Carbapenam substrate (**1**) is shown in light blue sticks, colored by atom type. An *F*_o_-*F*_c_ omit electron density map is shown for (**1**) (blue mesh, contoured at 3.0σ). Substrates, cofactors, and coproducts are shown in stick format. Distances between reactive groups are given in units of Å. (**C**) Product formation for the R280Q TokK variant. Substrate is shown in black spheres and the first methylated product is shown in pink squares. (**D**) Projections of the additional methyl groups added to C6 of (**1**) and their respective distances (in Å) from the 5’ carbon of 5’-dAH and the hydroxyl moiety of OHCbl, the latter of which serves as a surrogate of the active MeCbl cofactor. Spheres labeled Me and Et represent the suggested positions of the newly installed carbon atoms in the mono-methylated and dimethylated products.

The interactions between TokK and substrate **1** position the β-lactam appropriately both for activation of C6 by 5’-dA• and subsequent methyl addition by the Cbl cofactor (Figure 2D) (*17*). C6 of **1** is located directly in front of the 5’-carbon of 5’-dAH, 3.7 Å away, similar to other RS enzyme-substrate complexes that initiate reactions by H-atom abstraction. The orientation of these two groups in the structure does not reveal if the pro-*R* or pro-*S* H-atom is removed from C6 of **1** by 5’-dA•, as these substituents project equally above and below the 5’-carbon of 5’-dAH (Figure 2D). However, the structure does provide insight into the trajectory of methyl addition. C6 of **1** is located 4.2 Å above the axial ligand of Cbl (Figure 2D) at an angle of ∼85° relative to the 5’-carbon of 5’-dAH. This arrangement suggests that the methyl group adds to the bottom face of the β-lactam ring, consistent with the absolute configurations observed in the TokK products and thienamycin (*17, 18*). The distance and orientation of reactant functional groups in TokK also compares favorably to other enzymes that catalyze radical-mediated activation and functionalization of a substrate C-H bond, such as iron-dependent hydroxylases in the cytochrome P450 and 2-oxo-glutarate-(2OG)-dependent superfamilies (Figure S12). These systems orient their reactive groups similarly, but over a slightly shorter distance range (*19, 20*). This comparison highlights an important distinction between RS methylases and other radical functionalization enzymes. In P450s and Fe/2OG enzymes, a single reactive entity, a high-valent iron(IV)-oxo or iron(III)-hydroxo group, must both activate substrate and functionalize it. This strategy is inherently limiting because the enzyme can only activate and functionalize substrate from the same side. The Cbl-dependent RS radical functionalization platform is more versatile because the radical activation step is separated from methylation, allowing for more diverse stereochemical outcomes.

Intriguingly, the structure of TokK in complex with the substrate has marked structural similarities to another well-characterized RS methylase that does not rely on MeCbl, RlmN (Figure S13) (*21*). RlmN uses an *S*-methyl cysteinyl (methylCys) residue as an intermediate methyl carrier during the methylation of the *sp*^2^-hybridized C2 atoms of adenosine 2503 in ribosomal RNA and adenosine 37 in several transfer RNAs (tRNA). When the structure of RlmN crosslinked to an *E. coli* tRNA^Glu^ substrate is compared to that of substrate-bound TokK, the methylCys residue in the RlmN structure is in a position similar to that of the hydroxyl group of OHCbl in the TokK structure. Moreover, their respective substrates occupy similar positions in the active site (Figure S13). Although the catalytic mechanisms of these two enzymes are distinct, both obey a ping-pong kinetic model, wherein one SAM molecule is used to methylate the intermediate methyl carrier, while a second SAM molecule is used to generate a 5’-dA•.

Cbl is multifunctional in TokK, mediating both the polar methylation of Co(I) by SAM and the transfer of a methyl radical to C6 of the substrate. It is bound at the interface of the Cbl and RS domains with its dimethylbenzimidazole base (DMB) tucked into the Rossmann-fold of the N-terminal domain, a conformation termed “base-off” (Figure 3A **and** S6). OxsB and TsrM use a similar base-off approach to interact with their Cbl cofactors (Figure 3A **and** Figure S5), a binding mode that allows for extensive modulation of the reactivity of the Co(III) ion of MeCbl by the local protein environment (*22*). In TsrM, this structural feature is essential for the atypical polar methylation of its substrate, Trp, which requires heterolytic cleavage of the Co(III)-carbon bond of MeCbl. The bottom face of the Co ion in TsrM is adjacent to Arg69 but not directly coordinated, likely promoting nucleophilic attack of MeCbl by Trp by blocking coordination of a sixth ligand and destabilizing the Co(III)-C bond owing to charge-charge repulsion (Figure 3A) (*23–25*). In TokK, a different side chain, Trp76, occupies the lower axial face of the Cbl, residing 3.8 Å from the metal ion (Figure 3A). To investigate the role of this residue, Trp76 was substituted with Phe and Ala. Unexpectedly, rates of methylation slightly increased for both site-specific substitutions, while analysis by electron paramagnetic resonance (EPR) spectroscopy suggested that both variants exhibited the same 4-coordinate geometry as wild-type TokK (Figure S14). These data suggest that even when the steric bulk of Trp76 is reduced, water does not coordinate the Cbl. To perturb the local environment of the Cbl cofactor further, Trp76 was also substituted with His and Lys. The activity of Trp76His TokK resembles the activities of the Phe and Ala replacements, but the activity of Trp76Lys TokK was reduced by a factor of ∼50 for all three methylation steps (Figure 3B). The substitution tolerance of Trp76 in TokK contrasts with that of Arg69 of TsrM, which when substituted with Lys, was unable to transfer a methyl group from MeCbl to substrate (*11*). While Trp76 is not widely conserved among other well-characterized Cbl-dependent RS methylases, it is found in the same sequence context in the CysS primary structure (Figure S1).

**Figure 3.**
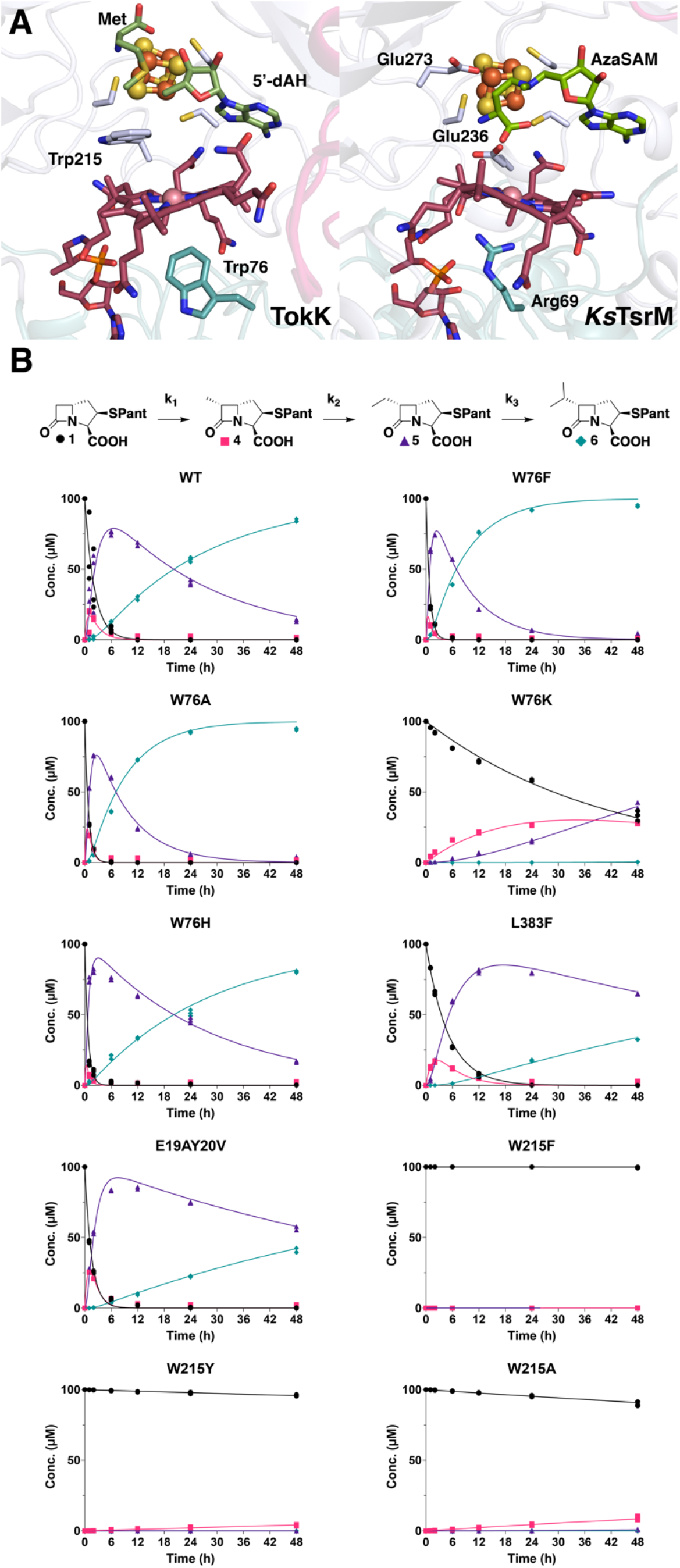
The Cbl and substrate binding sites influence overall activity and the relative rates of each TokK methylation step. (**A**) Comparative analysis of the side chains proximal to the Co ion in two cobalamin-binding RS enzymes, TokK and *Ks*TsrM (PDB ID: 6WTF). Selected amino acid side chains are shown in stick format and the Co ion is shown as a pink sphere. (**B**) Schematic diagram of the three sequential methylations performed by TokK. 48 h time course experiments performed in triplicate (each replicate represented as a symbol) tracking substrate (black spheres), methyl (pink squares), ethyl (purple triangles), and isopropyl (blue diamonds) product formation. Each product was estimated using COPASI (irreversible mass action model using the reaction scheme shown above) and simulated using Virtual Cell (shown as lines) for wild-type, W76F, W76A, W76K, W76H, L383F, E19AY20V, W215F, W215Y, and W215A.

The structure of **1** bound to TokK also rationalizes established differences in rate constants for each of the three methyl transfers catalyzed by this enzyme (Figure 3B). The second methylation to form the ethyl-containing carbapenam product **5** proceeds at least three-fold faster than the formation of **4**, a pattern that runs counter to known differences in reactivity of secondary and primary C-H bonds. Although we do not report a structure containing the singly-methylated intermediate **4**, if we presume that **4** remains anchored to Arg280, the methyl group at C6 would be positioned closer to the Cbl axial ligand and potentially at a more optimal angle than the original C6 C-H target. A third methylation to form the isopropyl carbapenem product **6** requires hydrogen atom abstraction from the same carbon, but the newly added ethyl carbon restricts the population of conformers accessible to 5’-dA•. This steric demand could help explain why k_3_ is markedly slower k_1_ and k_2_. The buried location of **1**, 5’-dAH, Met, and the Cbl cofactor suggests that dissociation of the methylated carbapenam products must occur prior to dissociation of the SAM cleavage products. This arrangement is consistent with the non-processive kinetic model used to fit the time-course kinetics of each TokK methylation. (*17*) A similar mechanism was proposed recently for CysS (Figure S1) (*6*). While CysS is only 29% identical both to TokK and to ThnK, all three proteins potentially contain a Trp side chain adjacent both to the Cbl and to the substrate (Figure S1). Substitution of Trp215 with Phe, Ala, or Tyr dramatically slows substrate methylation by TokK, suggesting it may play a role in catalysis (Figure 3B).

ThnK and TokK share 79.3% sequence identity and act on the same substrate, (2*R*)-pantetheinylated carbapenam (Figure S15), but ThnK performs two sequential methylations while TokK catalyzes three (*17, 26*). Nearly all of the residues in proximity to the active site are identical in the two orthologs. However, three non-conserved amino acids near the active site were explored to determine their role in controlling the extent of methylation. Leu383 is positioned deep in the active site and near 5’-dAH (Figure S15). When this residue is substituted with Phe, which is found at the same position in the primary structure of ThnK (Figure S15), the rate constants for all three methyl transfers are reduced (2.3-, 2.5-, and 4.5-fold for k_1,_ k_2_, and k_3_, respectively) compared to those of wild type TokK (Figure 3B). Two adjacent residues at the entrance to the pantetheine-binding tunnel, E19 and Y20, (Figure 2A **and** S15) were replaced with the cognate residues in ThnK to generate an E19A+Y20V double substitution. In this variant, the rate constant for the first methylation is increased 1.4-fold compared to that of wild type, and the rate constants for the second and third methylations are decreased 1.4- and 3.4-fold, respectively, therefore shifting the kinetic profile closer to the pattern observed with ThnK (k_1_ > k_2_, k_3_ = 0) (Figure 3B) (*17*).

The structure of TokK solved in the absence of substrate reveals remarkably few differences in overall fold or domain organization relative to the TokK•**1** complex (rmsd of 0.53 Å over 603 residues by C-alphas). The substrate binding channel, located at the interface of the three domains of TokK, remains intact without substrate with only modest alterations in size caused by the aforementioned conformational changes in the side chains of Arg280 and Tyr652 (Figure 2A **and** S9). The preformed nature of the substrate binding tunnel in TokK contrasts with observations from structures of TsrM in the absence and presence of its substrate, where a loop from the C-terminal moves to cap the active site in the presence of Trp (*11*). Although the C-terminal domain of TokK is significantly larger than that of TsrM, these domains appear to share a common role in substrate interaction (Figure S5).

The overwhelming majority of known Cbl-dependent RS methylases operate by the radical mechanism used by TokK, producing an equivalent of 5’-dAH and SAH for each methylated product molecule. The structure of the TokK active site reveals a scaffold for positioning the cofactors responsible for substrate activation and methyl transfer, and is consistent with the non-processive mechanism of sequential methylations observed for the enzyme, which requires the release of each partially alkylated intermediate and both SAM coproducts prior to reloading the active site for subsequent methylation. Moreover, the structure reveals that there is little active participation in catalysis from other amino acids in the active site, apart from substrate or cofactor binding. Interestingly, this scaffolding approach, which is underscored by the notable absence of conformation changes upon substrate binding, may be shared by the only other Cbl-dependent RS enzyme structurally characterized in complex with its substrate, TsrM. Although the electrophilic substitution mechanism used by TsrM requires a general base to accept the N1 proton of the indole ring of Trp during catalysis, biochemical studies and models of the functional enzyme-substrate complex suggest that the carboxylate group of a cosubstrate, SAM, functions in this capacity instead of an active site amino acid. Elucidation of additional structures and mechanisms for Cbl-dependent RS enzymes will reveal how a seemingly common approach for catalysis may be further elaborated within this large and diverse group of enzymes.

## Materials and Methods

### Overexpression and purification of TokK from Streptomyces tokunonesis

TokK (Uniprot ID: A0A6B9HEI0) was produced heterologously in *Escherichia coli* BL21 (DE3) via overexpression from a pET29b:*tokK*Tev construct (*17*). To facilitate production of soluble TokK protein with maximal occupancy of [4Fe–4S] cluster and cobalamin cofactors, the strain was transformed with two additional plasmids, pDB1282 and pBAD42-BtuCEDFB (*17, 27–29*). A 100 mL LB starter culture with 50 µg/mL kanamycin (pET29b) containing *tokK*), 50 µg/mL spectinomycin (pBAD42-BtuCEDFB), and 100 µg/mL ampicillin (pDB1282) was inoculated from a single colony and incubated for 18 h at 37 °C while shaking at 250 rpm. A 12 mL aliquot of the starter culture was used to inoculate a 4 L culture of LB medium supplemented with OHCbl (1.3 *µ*M) and allowed to shake at 180 rpm at 37 °C. Four of these cultures were grown to OD_600_ 0.3, at which point induction of genes on pDB1282 and pBAD42-BtuCEDFB was initiated with the addition of arabinose to a final concentration of 0.2%. To facilitate iron-sulfur cluster incorporation, 25 *µ*M FeCl_3_ and 150 *µ*M cysteine were added at the initiation of induction with arabinose. The cultures were then grown to OD_600_ 0.6 followed by incubation in an ice–water bath for 1 h. IPTG was added to a final concentration of 1 mM, and the cultures were incubated at 18 °C for an additional 18 h before harvesting the cells by centrifugation at 6000 × *g*. The resulting cell paste (∼50 g) was flash frozen in liquid N_2_ and stored in liquid N_2_ prior to protein purification.

All purification steps and subsequent manipulations of TokK were performed in a Coy Laboratory Products vinyl anaerobic chamber. Cell paste was resuspended in 100 mL of lysis buffer (50 mM HEPES, pH 7.5, 300 mM KCl, 10% glycerol, 5 mM imidazole, 10 mM BME). The cell suspension was incubated with lysozyme (1 mg/mL), DNase (0.1 mg/mL), and PMSF (0.18 mg/mL) for 30 min at room temperature and then cooled to 4 °C (*17, 29*). Cells were then lysed by sonic disruption (70% amplitude, 45 s on, 59 s off, ∼15 min) and then centrifuged at 50000 × *g* for 1 h to separate insoluble material. The resulting supernatant was loaded onto a pre-equilibrated column of Ni-NTA resin and purified by immobilized-metal affinity chromatography. The protein-loaded resin was washed with 50 mL of lysis buffer prior to addition of elution buffer (lysis buffer supplemented with 500 mM imidazole). Dark colored elution fractions were pooled and concentrated to 1 mL in an Amicon 10 kDa MWCO ultrafiltration device (EMD Millipore; Billerica, MA). The protein fractions were then exchanged into TEV protease cleavage buffer (50 mM HEPES, pH 7.5, 300 mM KCl, 15% glycerol, 10 mM BME) using a PD-10 pre-poured gel-filtration column from GE Biosciences (Pittsburgh, PA). The exchanged protein sample was allowed to react with 10 units of TEV protease (∼2 mg/mL) in 25 mM Tris-HCl pH 8.0, 50 mM NaCl, 1 mM TCEP, and 50% glycerol (Millipore Sigma; St Louis, MO) for two days on ice to generate TokK containing only 6 additional amino acids (ENLYFQ) on its N-terminus. On the second day, the [4Fe–4S] cluster and cobalamin cofactors were reconstituted as previously described (*29*). The reaction mixture was then reapplied to the Ni-NTA resin to capture any remaining His-tagged protein, and purified ENLYFQ-TokK was collected in the flow-through fraction. TokK was concentrated and exchanged into gel-filtration buffer (50 mM HEPES, pH 7.5, 300 mM KCl, 1 mM DTT, 15% glycerol) for size-exclusion chromatography. In this step, the protein was applied to a HiPrep 16/60 S200 column using an ÅKTA fast protein liquid chromatography (FPLC) system (GE Biosciences) housed in the anaerobic chamber. TokK elutes as an apparent monomer. Fractions were pooled based on UV-vis absorption at 280 and 410 nm and concentrated to 38 mg/mL.

### X-ray crystal structure determination of TokK

#### General crystallographic methods

X-ray diffraction datasets were collected at the General Medical Sciences and Cancer Institutes Collaborative Access Team (GM/CA-CAT) and at the Advanced Photon Source, Argonne National Laboratory and Berkeley Center for Structural Biology (BCSB) beamlines at the Advanced Light Source at Lawrence Berkeley National Laboratory. All datasets were processed using the HKL2000 or HKL3000 package, and structures were determined by single anomalous dispersion (SAD) phasing using Autosol/HySS or by molecular replacement using the program PHASER (*30–33*). Model building and refinement were performed with Coot and phenix.refine (*30, 34*). Figures were prepared using PyMOL (*35, 36*). Substrate channel figures were prepared using Hollow (*37*). Ligplot was utilized to visualize the binding of substrate 1 (*38*).

#### Crystallization and structure solution of 5’dAH•Met•TokK

Purified TokK protein aliquots were diluted to 8 mg/mL in 34 mM HEPES, pH 7.5. 5’-dAH and Met (Millipore Sigma, St. Louis, MO) were added to the resulting solution to final concentrations of 2 mM each. The mixture was incubated for 30 min at room temperature. In hanging-drop vapor-diffusion trials with 100 mM magnesium chloride, 100 mM calcium chloride, 20% PEG 8000, and 10% 1,6-hexanediol as the precipitating reagent, brown plate-shaped crystals appeared within three days. Trials were initiated by adding 1 µl of protein with 1 µl precipitating solution, followed by equilibration against 500 uL of a 0.5 M LiCl well solution at room temperature. Crystals were prepared for data collection by mounting on rayon loops followed by brief soak in cryoprotectant solution (50% (*v/v*) precipitating reagent and 50% (*v/v*) ethylene glycol) and flash-freezing in liquid nitrogen.

Diffraction datasets for single-wavelength anomalous diffraction phasing were collected at the iron K-edge x-ray absorption peak (λ = 1.72194 Å) with 360° of data measured using a 0.5° oscillation range to 2.52 Å resolution. Additionally, a 1.79-Å resolution native dataset was collected at λ = 1.03313 Å (Table S1). Heavy atom sites were identified using HySS implemented within Phenix Autosol (*30*). The initial overall figure-of-merit (FOM) was 0.265 and the Bayes CC was 18.2 (*30*). Phenix Autobuild was used to generate an initial model of 541 residues out of 687 in chain A and 569 residues out of 687 in chain B with R_work_/R_free_ of 0.25/0.31. Iterative manual model building and refinement were performed in Coot and Phenix (*34*). This model was used to obtain phase information for the 1.79-Å resolution native dataset by molecular replacement using Phenix Phaser-MR (*30*). Geometric restraints for 5’-dAH and cobalamin were obtained from the Grade Web Server (Global Phasing). *R_free_* flags were maintained so that the same 5% of the reflections were used as the test set. The final model consists of residues 9-412, 416-672, one [4Fe-4S] cluster, one hydroxycobalamin cofactor, one methionine (Met), and one 5’-deoxyadenosine (5’-dAH) in Chain A; residues 7-413, 416-672, one [4Fe-4S] cluster, one hydroxycobalamin cofactor, one Met, and one 5’-dAH in Chain B. The final model also contains two chloride ions, two potassium ions, 21 molecules of ethylene glycol, and 887 waters. The Ramachandran plot shows that 97.4% residues are in favored regions with the remaining 2.6% in allowed regions (*35*). Data collection and refinement statistics are shown in Table S1.

#### Synthesis of substrate

Synthesis of the TokK substrate, (2*R*,3*R*,5*R*)-3-((2-(3-((*R*)-2,4-dihydroxy-3,3-dimethylbutanamido)propanamido)ethyl)thio)-7-oxo-1-azabicyclo[3.2.0]heptane-2-carboxylic acid, was carried out as previously described (*26*). Characterization matched that previously reported.

#### Crystallization and structure solution of substrate-bound 5’dAH•Met•TokK

Purified TokK was diluted to 8 mg/mL TokK in 34 mM HEPES, pH 7.5. 5’-dAH, Met, and substrate were added to final concentrations of 2 mM each. The solution was incubated for 15 min at room temperature. In hanging drop vapor diffusion crystallization trials with 0.2 M lithium sulfate, 0.1 M Tris–HCl, pH 8.5, and 18% PEG 8000 as the precipitant, brown plate-shaped crystals appeared within two weeks. Trials were initiated by mixing 1 µl of protein and 1 µl of precipitant followed by equilibration at room temperature against a 500 uL reservoir of the precipitating solution. Crystals were prepared for data collection by mounting on rayon loops followed by a brief soak in perfluoropolyether oil (Hampton Research) for cryoprotection and flash-freezing in liquid nitrogen.

The structure containing substrate was solved by molecular replacement (Phaser-MR in Phenix) using the coordinates of the 5’dAH•Met•TokK as the search model. Iterative manual model building and refinement were performed in Coot and Phenix (*30, 34*). Initial geometric restraints for (2*R*)-pantetheinylated carbapenem substrate were generated by eLBOW in Phenix (*30*). Final geometric restraints for the carbapenem substrate were created by using the PRODRG 2 server (*39*). The final model consists of residues 8-566, 571-672, one [4Fe-4S] cluster, one hydroxycobalamin cofactor, one Met, one 5’-deoxyadenosine (5’-dAH), and one (2*R*)-pantetheinylated carbapenem substrate in chain A; residues 9-672, one [4Fe-4S] cluster, one hydroxycobalamin cofactor, one Met, and one 5’-dAH in chain B. The final structure also contains two glycerol molecules, four potassium ions, and 1193 waters. The Ramachandran plot showed that 97.7% of residues are in favored regions with the remaining 2.3% in allowed regions (*35*). Data collection and refinement statistics are shown in Table S1.

#### Cobalamin ligand assignment in protein crystals

The protocol was adapted from previously published studies (*27*). Approximately 15 TokK + 5’-dAH + Met crystals were looped from the crystallization drop into 40 uL of mother solution in a darkly colored Eppendorf tube. In the dark, 50 mM H_2_SO_4_ was added to the tube. The tube was vortexed and then centrifuged for 20 min. 5 µl of this solution was injected into a Thermo Fisher Scientific UHPLC/QExactive HF-X mass spectrometer equipped with a C18 column (2.1 × 100 mm) equilibrated in 5% solvent A (0.1% formic acid) and 95% solvent B (0.1% formic acid in acetonitrile). The solvent B composition was increased to 98% from 1 to 7 min. Cobalamin forms were detected by ESI^+^, scanning from *m/z* 150-1700 with a resolution of 120,000. A calibration curve (0.1 µM – 5 µM) of cobalamin standards was run concurrently to quantify the cobalamin forms in the sample.

#### Generation of sequence similarity networks (SSNs)

A pool of annotated Cbl-dependent radical SAM enzymes was generated by merging representative annotated B_12_-dependent radical SAM enzyme sequences from radicalsam.org (megacluster 2-1, subgroup 5, https://radicalsam.org/explore.php?id=cluster-2-1&v=3.0, accessed March 2021) with those from the USCF structure-function linkage database (SFLD) (http://sfld.rbvi.ucsf.edu/archive/django/index.html, accessed April 2021) (*40, 41*). Additional sequences were included for the following functionally characterized enzymes (with Uniprot accession codes): *Sl*TsrM (C0JRZ9), *Ks*TsrM (E4N8S5), GenK (Q70KE5), GenD1 (Q2MG55), PhpK (A0A0M3N271), Fom3 (Q56184), CysS (A0A0H4NV78), OxsB (O24770), PoyC (J9ZXD6), swb7 (D2KTX6), ArgMe (Q8THG6), ThnK (F8JND9), TokK (A0A6B9HEI0), BchE (Q7X2C7), CouN6 (A0A1H2F7M3), and CloN6 (Q9F8U1) (*6, 11, 13, 14, 17, 42-50*). The pool of B_12_-dependent radical SAM enzyme sequences from radicalsam.org contains 9,724 representative entries reflecting 53,470 unique sequences. The class B and B_12_-binding radical SAM enzyme sequence groups from the SFLD contain 1,525 and 5,920 representatives, respectively. Each group reflects a total pool of 4,232 and 15,983 unique sequences, respectively (*40*). The final represent sequence set used for SSN generation contained approximately 11,000 sequences after removal of duplicates. This dataset represents a larger sequence pool of >50,000 unique entries.

The Enzyme Function Initiative enzyme similarity tool (EFI-EST) (https://efi.igb.illinois.edu) was used to perform an all-by-all BLAST analysis of the representative sequence dataset described above to create an initial SSN (*41, 51–53*) with an alignment score threshold of 65. To eliminate protein fragments, sequence length was restricted to greater than 300 amino acids. The final SSN (Figure 1C **and** S2) is depicted as a representative node network in which each node reflects sequences with >40% identity. All networks were visualized with the Organic layout in Cytoscape (*54*). The SSN in Figure 1C represents 36 sequence clusters extracted from the full SSN shown in Figure S2. The clusters in Figure 1C were selected based on number of nodes or the presence of functionally annotated or structurally characterized sequences.

#### Preparation of EPR Samples

TokK was diluted to a final concentration of approximately 250 µM for each EPR sample. The samples were photolyzed on ice for 45 minutes to generate the cob(II)alamin state. All samples were flash-frozen in cryogenic liquid isopentane in an anaerobic chamber. The resulting samples were stored in liquid nitrogen prior to analysis. EPR measurements were taken on a Magnettech 5000 x-band ESR spectrometer equipped with an ER 4102ST resonator. Temperature was controlled by an ER 4112-HV Oxford Instruments (Concord, MA) variable-temperature helium-flow cryostat. All measurements were taken at 70K, with 1mT modulation amplitude, and 1mW power.

#### Construction of TokK Variants

TokK variants were generated by overlap extension PCR using pET29b:*tokK*Tev as a template and primers described in Table S2. TokK_F_s was used as the forward primer for all constructs except the E19A/Y20V double variant, for which TokK_F_EYmut was used instead. After amplification, PCR products were digested with *Nde*I and *Xho*I and ligated into a similarly digested pET29b vector. Sequence-verified constructs were used to transform *E. coli* BL21 (DE3) along with helper plasmids pDB1282 and pBAD42-BtuCEDFB as described above for overexpression.

#### Methylation Assays of TokK Variants

Expression and purification of TokK variants was carried out as previously described for wild-type TokK (*17*), except reconstitution (*29*) was done concurrently with overnight TEV protease cleavage, and an additional buffer exchange was done using an Econo-Pac 10DG column (Bio-Rad) to remove excess reconstitution reagents. Methylation assays were carried out in triplicate and contained 100 mM HEPES, pH 7.5, 200 mM KCl, 1 mM SAM, 1 mM methyl viologen, 2 mM NADPH, 0.5 mM methylcobalamin, 100 μM substrate, and 100 μM enzyme. At each time point, an aliquot of the reaction mixture was diluted 5×, filtered through a 10 kDa Amicon ultrafiltration device, and analyzed for product formation using UPLC-HRMS as previously described (*17*). First order rate constants were determined using the COPASI parameter estimation tool (*55*), and curves were simulated using Vcell (*56*).

## Supplementary Information

**Figure S1.**
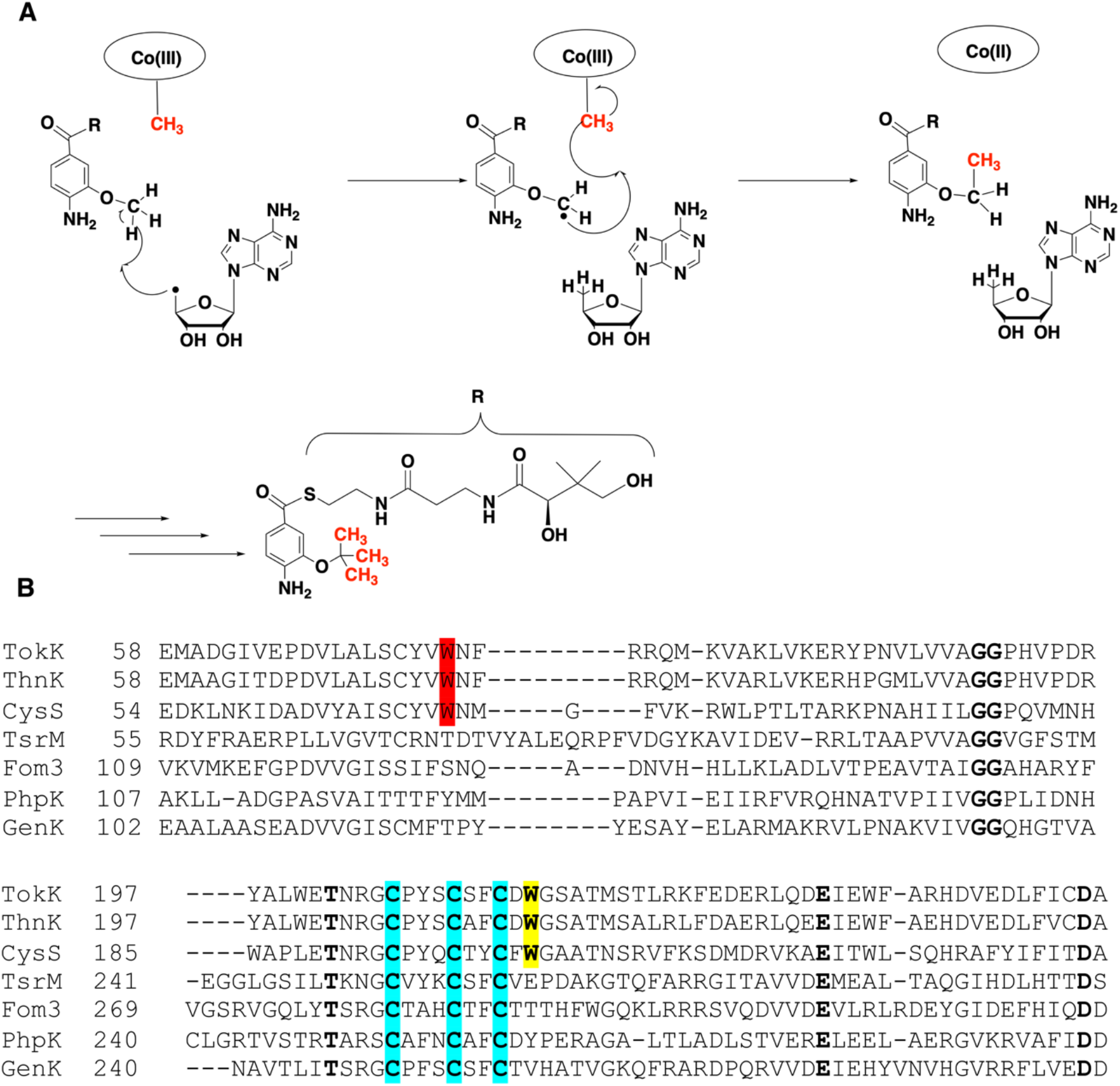
Comparison to CysS, another class B sequential methylase. CysS performs sequential radical methylations like TokK and ThnK (**A**). Partial alignment of TokK (Uniprot ID: A0A6B9HEI0), ThnK (Uniprot ID: F8JND9), CysS (Uniprot ID: A0A0H4NV78), TsrM (Uniprot ID: C0JRZ9), Fom3 (Uniprot ID: Q56184), PhpK (Uniprot ID: A0A0M3N271), and GenK (Uniprot ID: Q70KE5) (**B**). Cysteines that coordinate the iron-sulfur cluster are highlighted in blue. Trp76 (highlighted in red) and Trp215 (highlighted in yellow) in TokK is conserved in ThnK and CysS, but not in other known cobalamin-binding RS methylases. Completely conserved residues are bolded.

**Figure S2.**
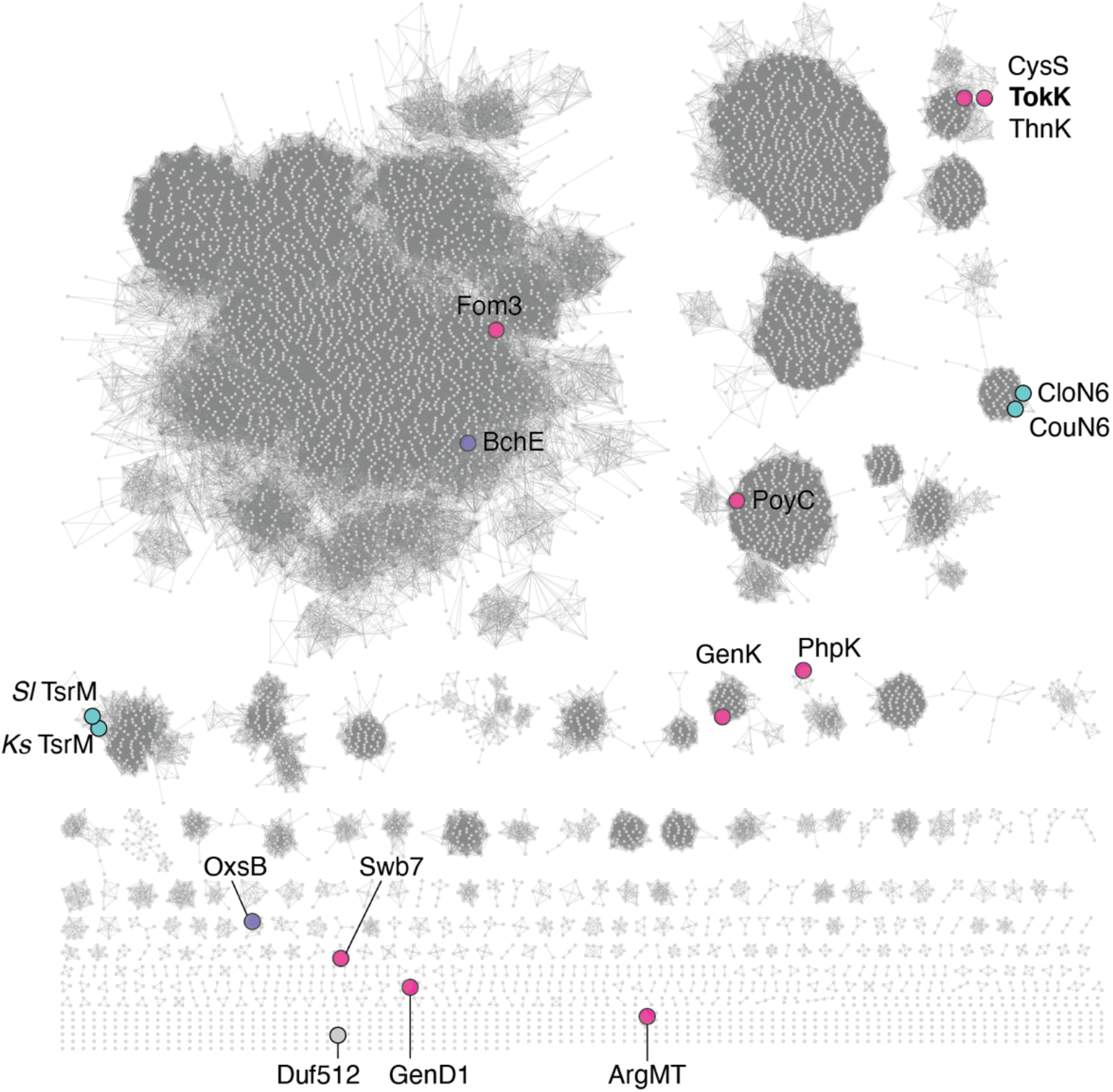
A sequence similarity network (SSN) of Cbl-dependent RS enzymes. An SSN of ∼11,000 sequences of Cbl-dependent RS enzymes indicates that they group into many distinct clusters. Sequences were compiled with an alignment score of 65. Each node represents either a single sequence or a set of sequences with >40% sequence identity. Enlarged and colored nodes correspond to sequences with proposed or validated functions (coloring as shown in Fig. 1C: pink = radical methylases, purple = non methylases, teal = polar methylases, gray = unknown function). Duf512 and ArgMT are structurally distinct RS enzymes that lack a traditional Rossmann fold Cbl-binding domain and were included in this analysis as outgroup sequences. As such, they are found among the singletons in the network.

**Figure S3.**
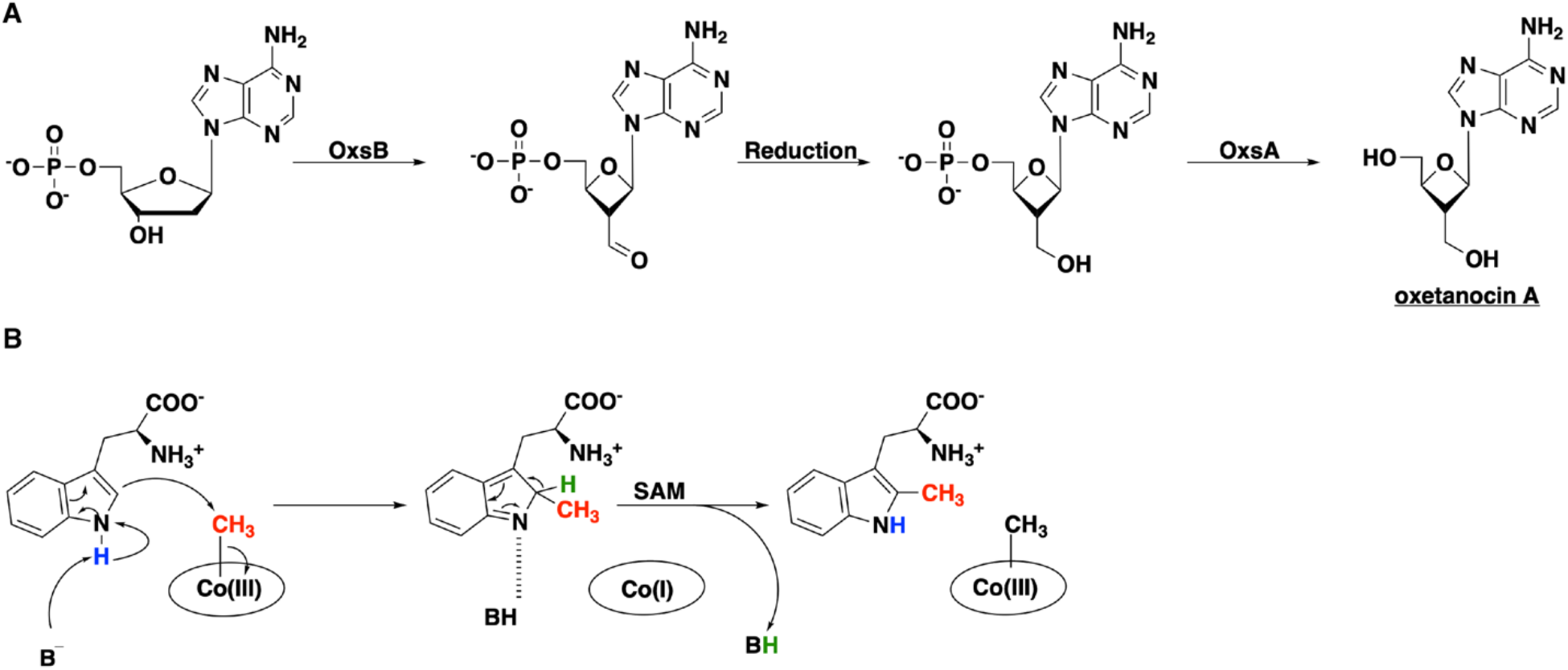
Reactions performed by OxsB and TsrM, notable Cbl-binding radical SAM enzymes. (**A**) Proposed pathway for the biosynthesis of oxetanocin A by OxsB and OxsA. Currently the aldehyde reduction step is unknown. (**B**) Proposed non-radical reaction performed by TsrM. The carboxylate in SAM is implicated in playing a dual role in catalysis, both as the source of the methyl donor and as the base to prime the substrate for nucleophilic attack.

**Figure S4.**
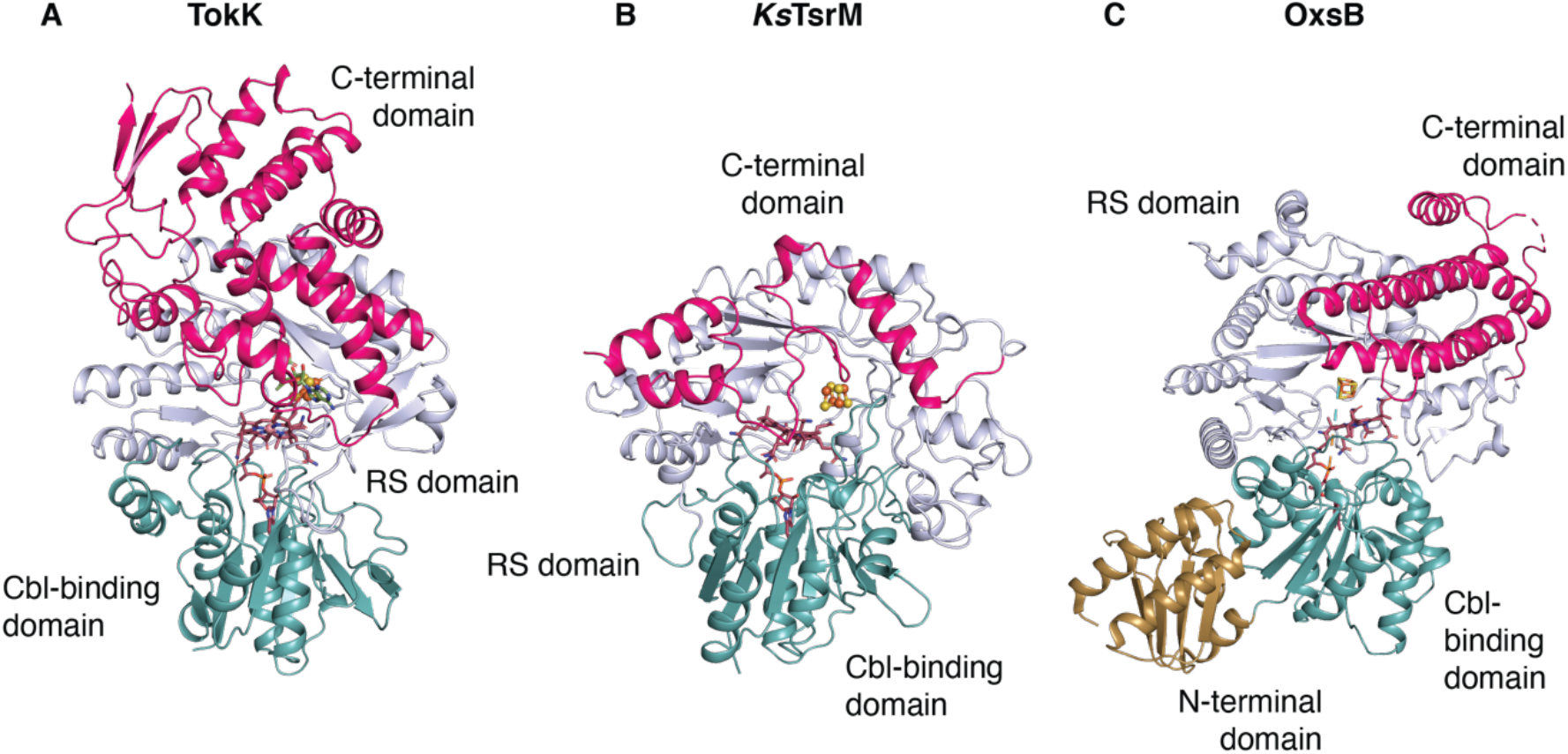
Structural comparisons of Cbl-dependent RS enzymes. The overall structures of TokK (A), *Ks*TsrM (PDB ID: 6WTE) (B), and OxsB (PDB ID: 5UL3) (C) are shown as ribbon diagrams and colored by domain (Cbl-binding domain, teal; RS domain, light blue; and C-terminal domain, pink). OxsB has a fourth domain, an N-terminal domain of an unknown function, shown in gold. All three enzymes share very similar Cbl-binding domains with a characteristic Rossmann fold. However, as shown in Fig. S5, the RS domain and *C*-terminal domain differ in each system.

**Figure S5.**
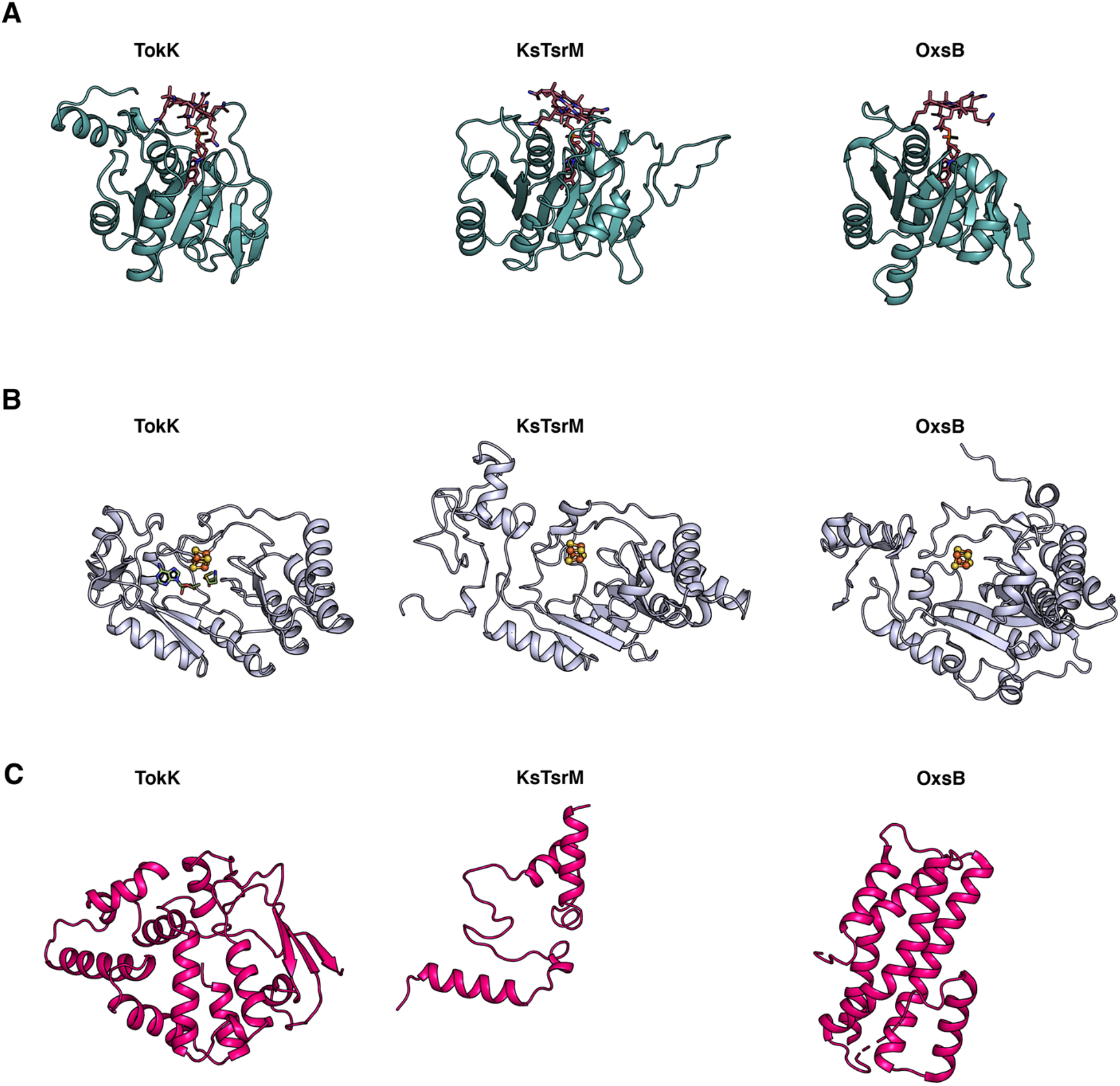
Comparison of the Cbl-binding, RS, and C-terminal domains of TokK, *Ks*TsrM, and OxsB. The domains are colored as in fig. S3. (**A**) The Rossmann fold, in teal, is highly similar among TokK, *Ks*TsrM, and OxsB. (**B**) The core of each of the RS domains is a (β/⍺)_6_ motif; however, there are distinct differences. The RS domain of OxsB is more compact than those of TokK or *Ks*TsrM, and all three have unique extra secondary structure features. (**C**) The C-terminal domains for TokK, *Ks*TsrM, and OxsB are vastly different in architecture.

**Figure S6.**
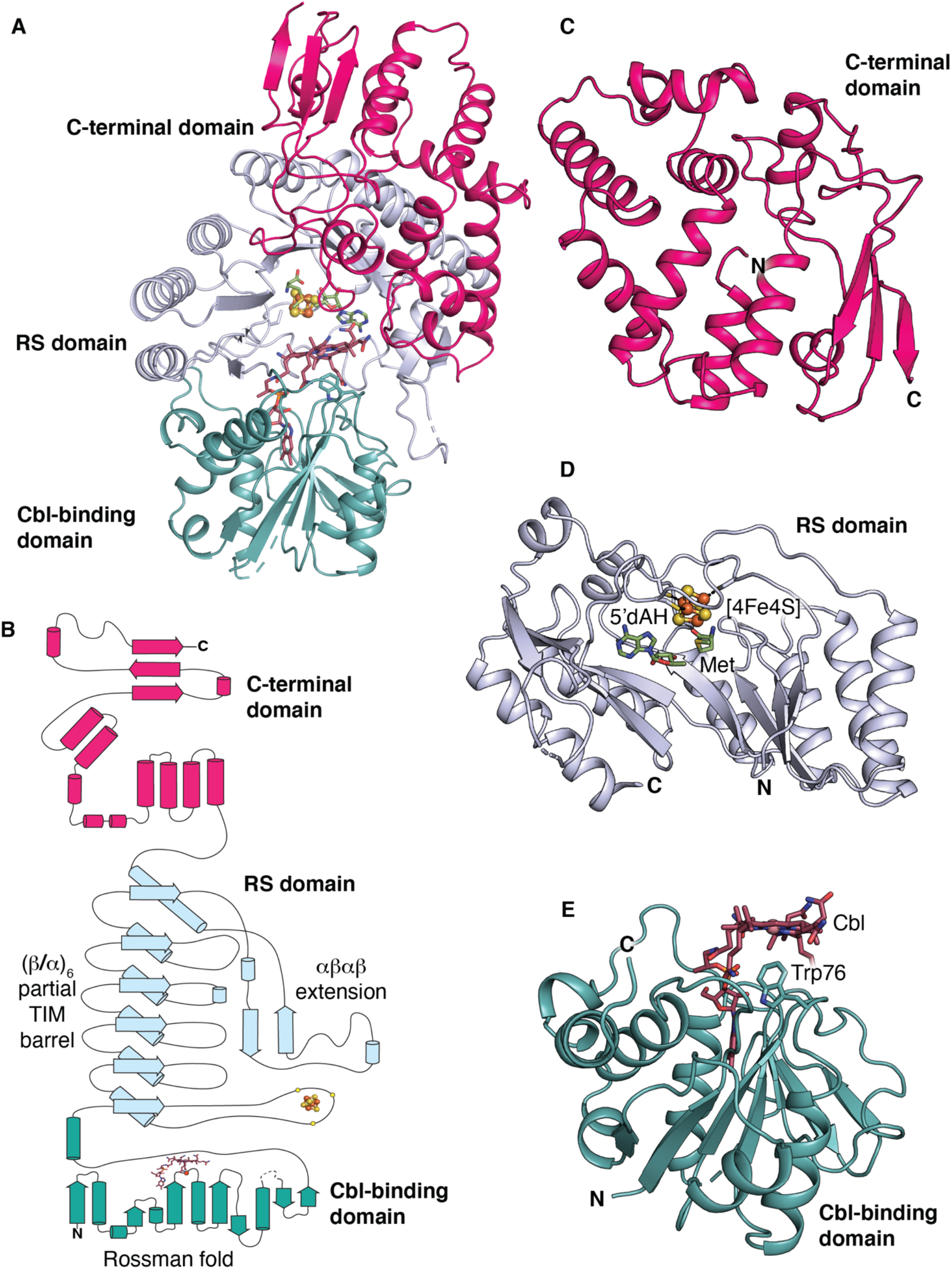
Domain architecture of TokK. (**A**) The three domains of TokK are portrayed in a ribbon diagram. The N-terminal Cbl-binding domain is shown in teal, the RS domain is shown in light blue, and the C-terminal domain is shown in pink. (**B**) A topology diagram of TokK with domains colored as in panel A. The cobalamin is portrayed in sticks, and the lower axial Trp side chain is shown as a red dot. The iron-sulfur cluster is shown in orange and yellow spheres, and the ligating Cys residues are shown as small yellow spheres. Zoomed in views of the C-terminal domain (**C**), RS domain (**D**), and Cbl-binding domain (**E**) are shown as ribbon diagrams.

**Figure S7.**
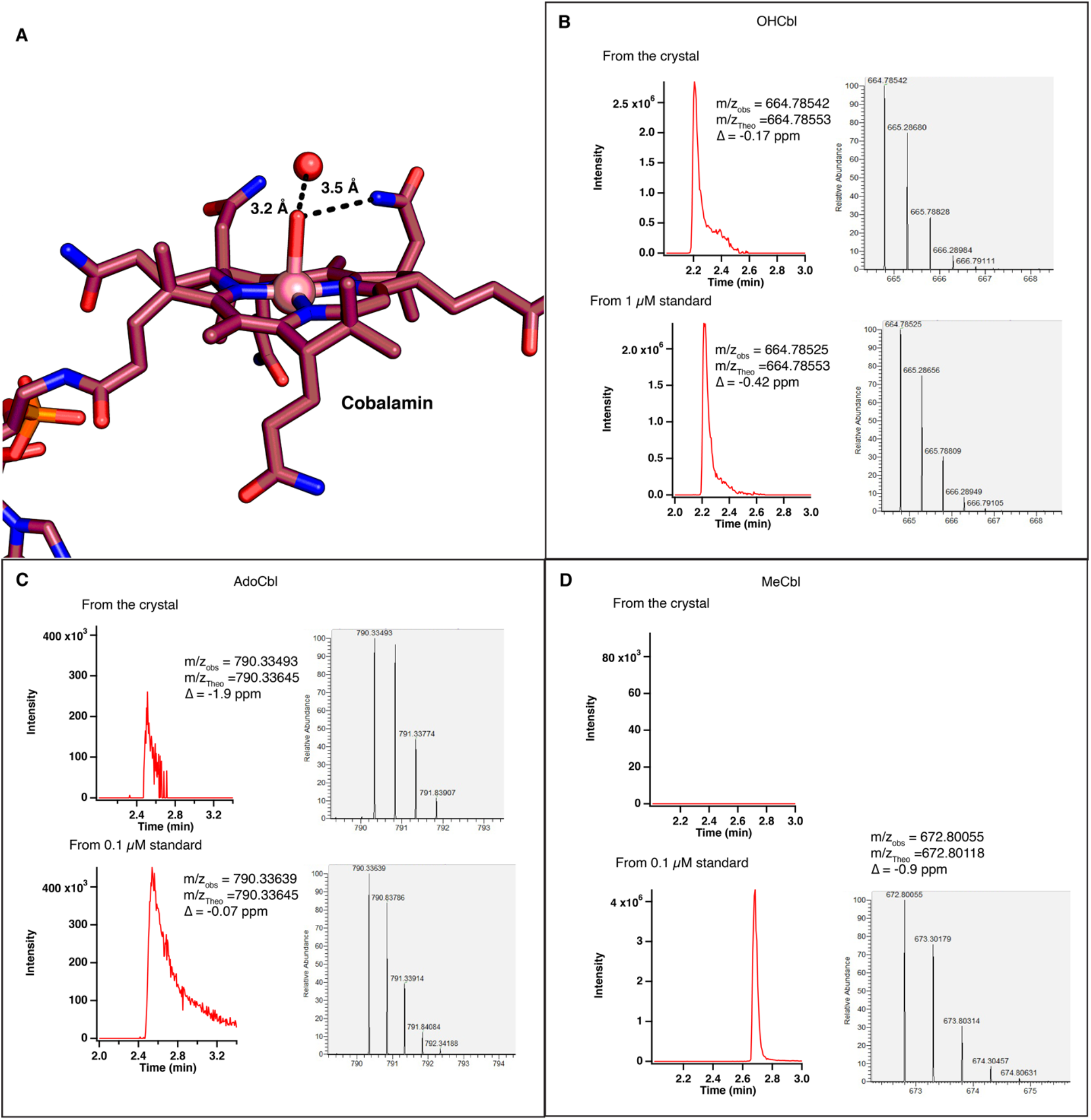
Assignment of the Cbl top ligand in structures of TokK crystals. (**A**) Hydrogen bonds to the top axial ligand above the cobalamin, suggesting a hydroxyl group. (**B**) Mass spectrometric analysis of OHCbl content in TokK + 5’-dAH + Met crystals with comparison to a 1 µM OhCbl standard, with relative abundance of the masses observed. (**C**) Mass spec analysis of AdoCbl content in TokK + 5’-dAH + Met crystals with comparison to a 0.1 µM AdoCbl standard, with relative abundance of the masses observed. (**D**) Mass spec analysis of MeCbl content in TokK + 5’-dAH + Met crystals with comparison to a 0.1 µM MeCbl standard, with relative abundance of the masses observed. As can be seen, the crystals are composed almost exclusively of OHCbl.

**Figure S8.**
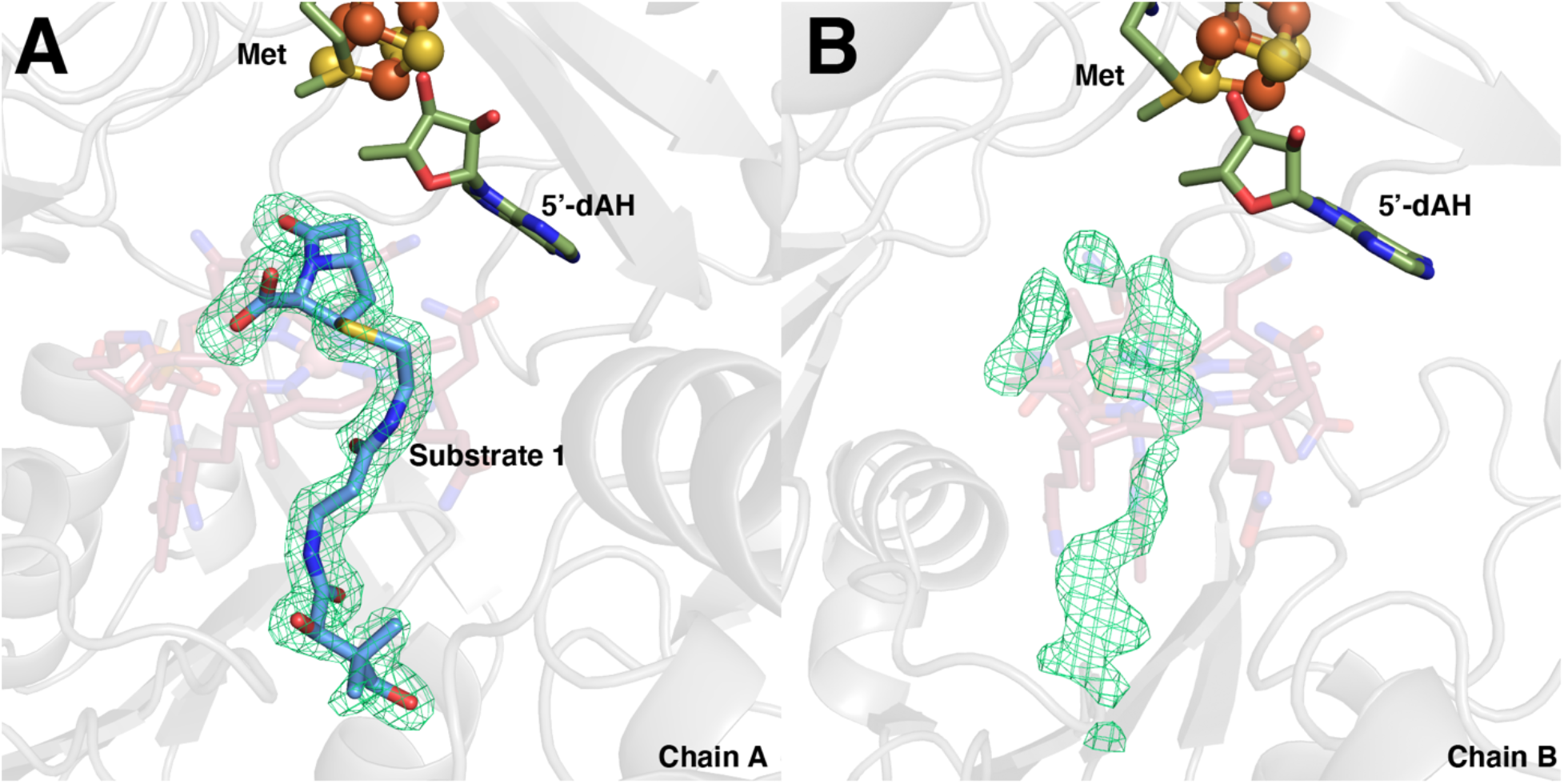
Comparison of the substrate-binding site in each subunit of the TokK•substrate complex. Omit *F*_o_-*F*_c_ electron density maps (green mesh, contoured at 3.0σ) shown for chain A (**A**) and chain B (**B**).

**Figure S9.**
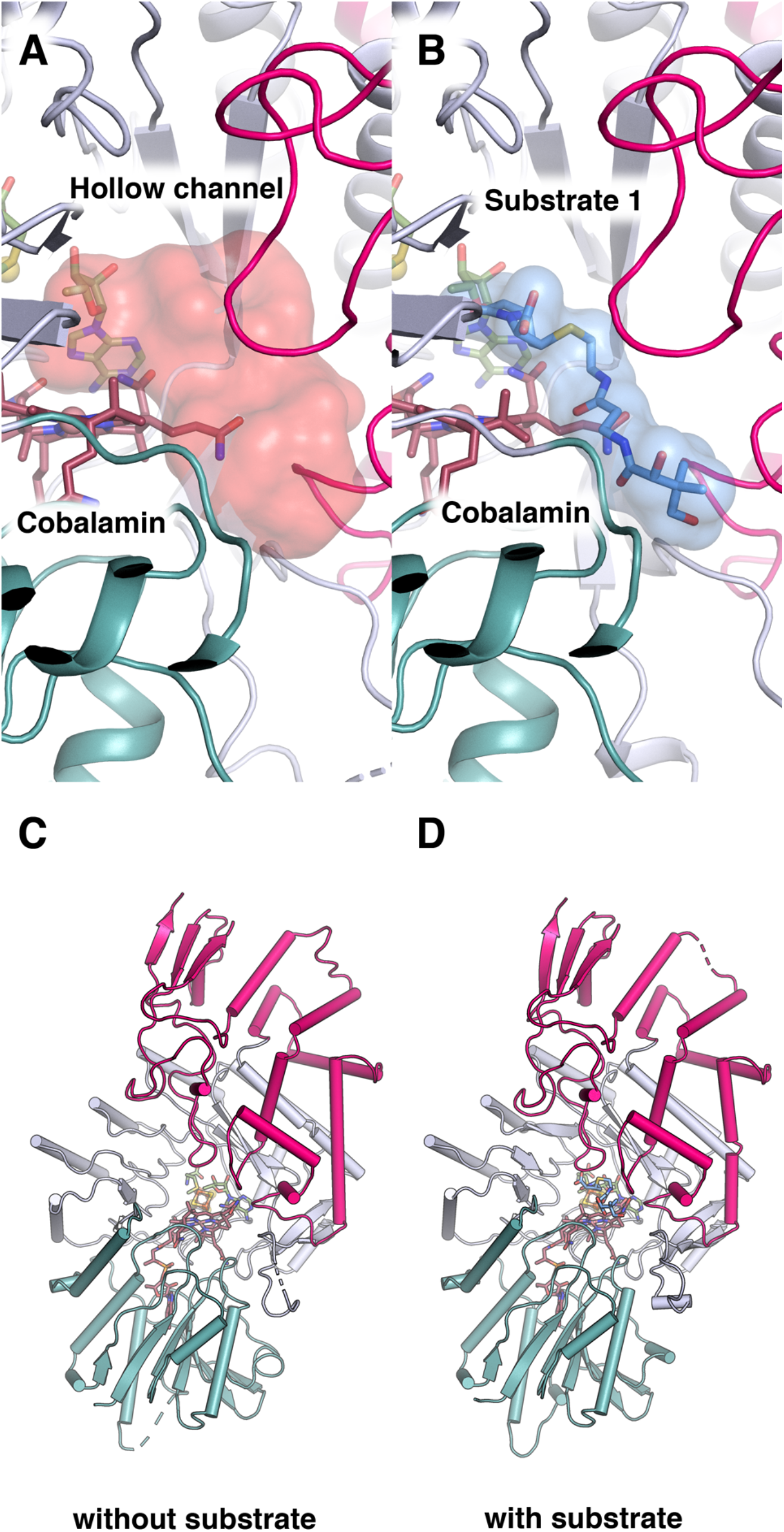
Comparison of TokK structures with and without substrate. (**A**) HOLLOW model shown in red, indicating the substrate channel of 5’-dAH and Met-bound TokK at the interface of the Rossmann fold (teal) and C-terminal (pink) domains. The channel is approximately 22 Å long and 15 Å wide. (**B**) Binding of the substrate of TokK is shown in light blue. The surface of the substrate is displayed in light blue, showing how the substrate binds in the channel predicted by HOLLOW. The substrate, approximately 16 Å long and 5 Å wide at the ß-lactam ring, is easily accommodated by the channel in the TokK structure in the absence of substrate. (**C**) Overall structure of TokK in the absence of substrate, with domains colored accordingly. (**D**) Overall structure of substrate-bound TokK. There are very few overall structural fold differences between the two structures (r.m.s.d. of 0.579 Å over 603 residues).

**Figure S10.**
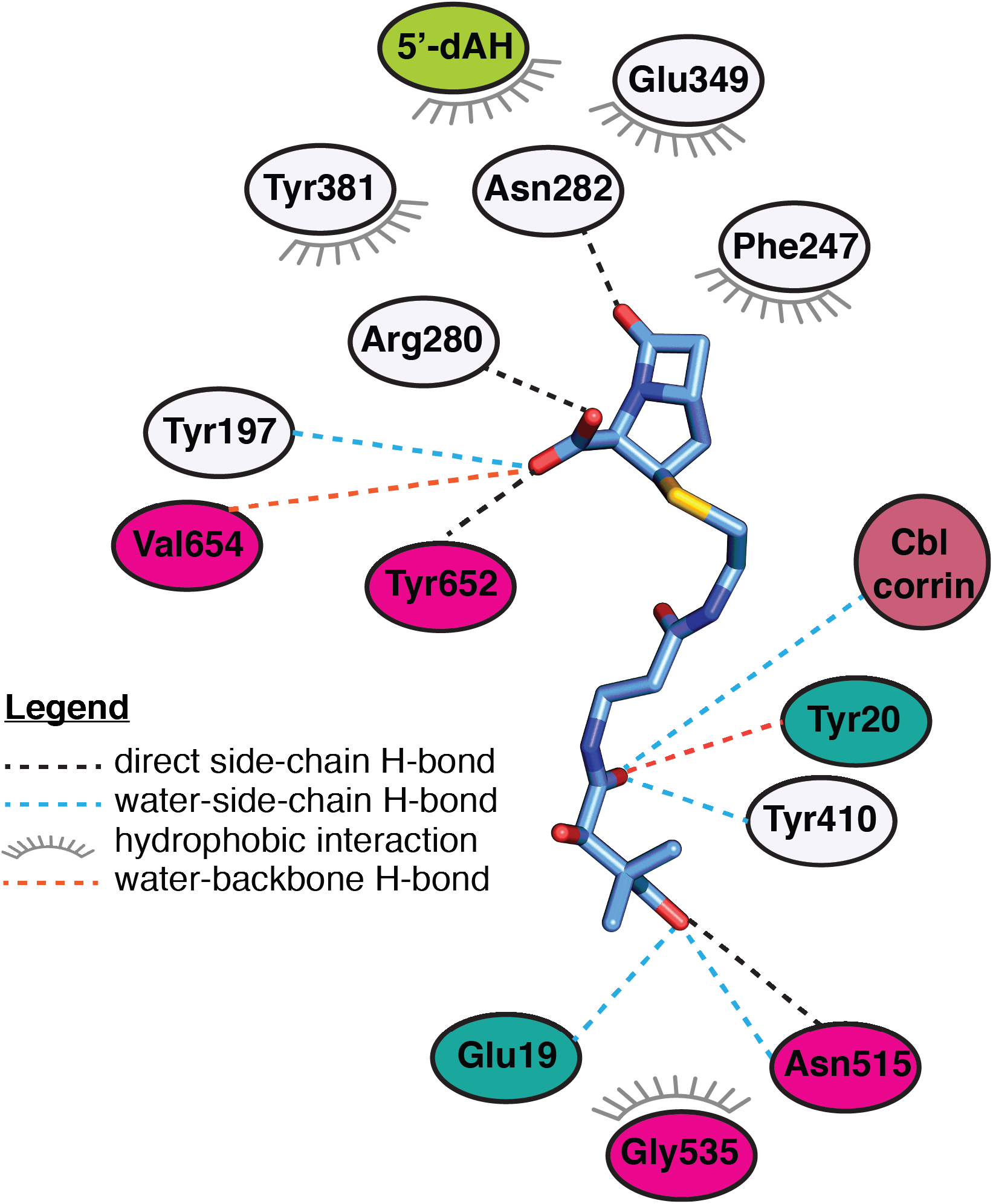
Diagram of carbapenam interactions in the TokK complex with substrate. Carbapenam substrate **1** shown in blue sticks and colored by atom type. Polar and hydrophobic interactions were mapped with the LigPlot program and indicated with dashed lines or a starburst symbol.

**Figure S11.**
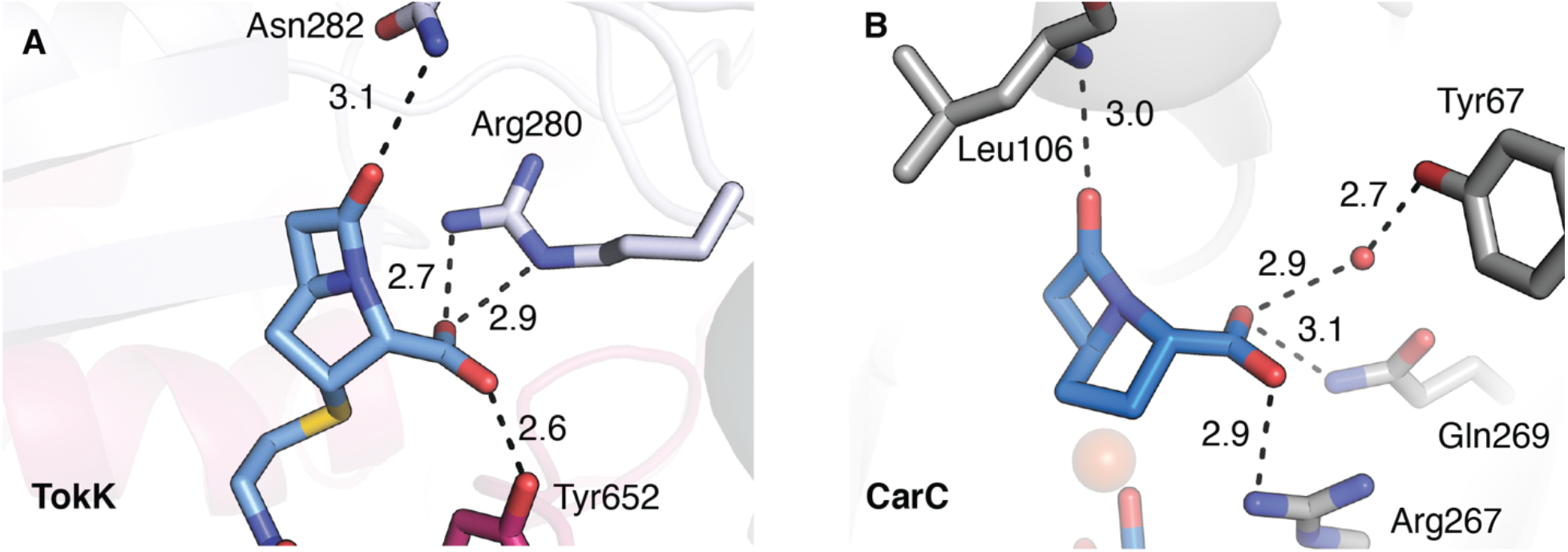
Comparison of ß-lactam binding sites in TokK and the β-lactam epimerase and desaturase, CarC. Residues that interact with the polar substituents in the β-lactam ring are shown in stick format for TokK (**A**) and CarC (PDB ID: 4OJ8) (**B**). While the two enzymes are structurally distinct, they use a similar number and type of functional groups to anchor the β-lactam by using direct and water-mediated contacts to the C7 carbonyl and C3 carboxylate substituents.

**Figure S12.**
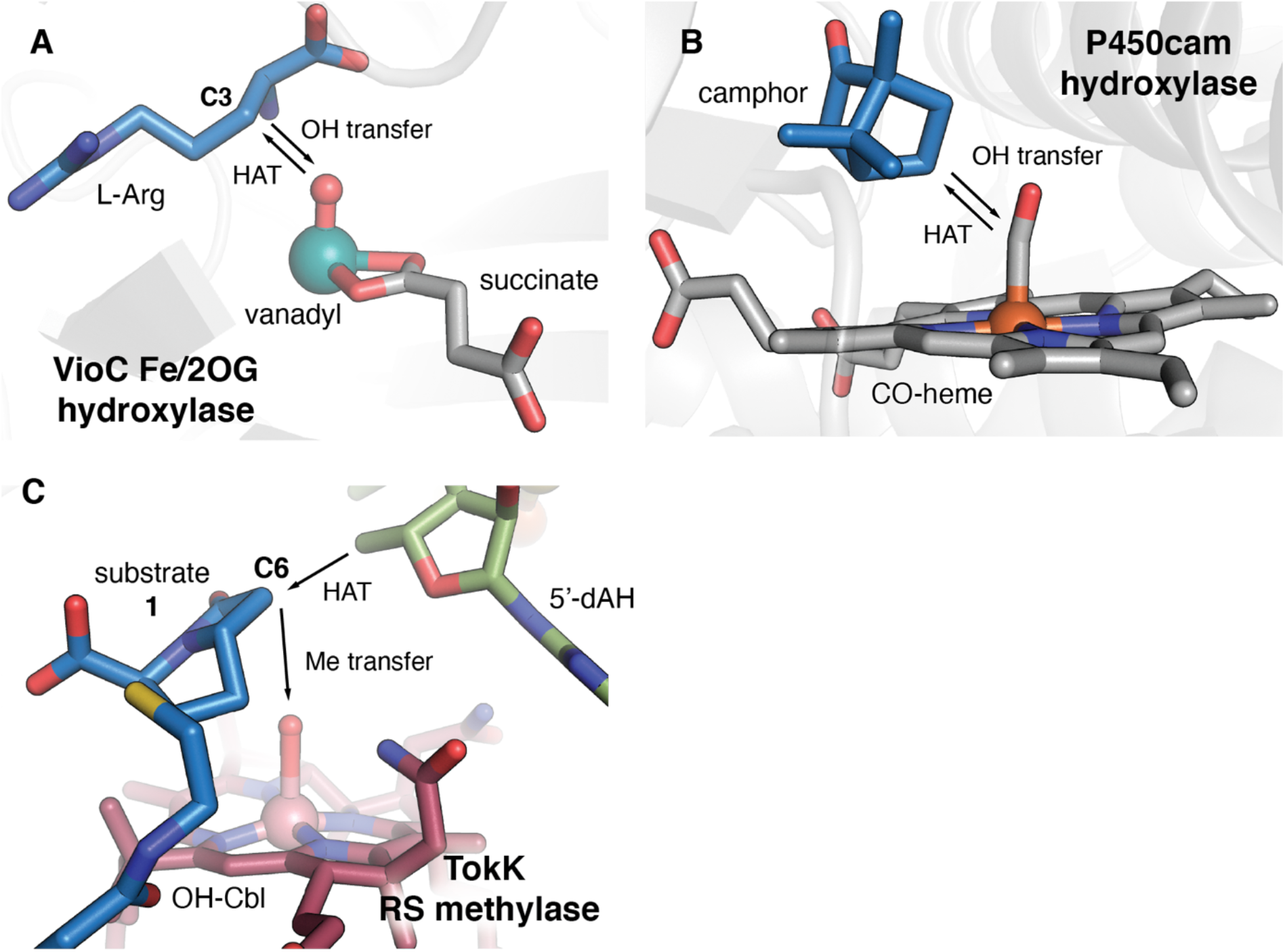
A comparison of the active sites of radical-mediated hydroxylase and methylase active sites. This analysis reveals differences in the orientation of hydrogen atom transfer (HAT) intermediates and -OH or -CH_3_ functionalization moieties. Fe(II)- and 2-oxo-glutarate-dependent (Fe/2OG) oxygenases use a ferryl [Fe(IV)-oxo)] intermediate to abstract an H-atom from an unactivated substrate carbon. The resulting Fe(III)-OH complex then couples with the substrate radical to yield a hydroxylated product. A vanadyl [V(IV)-oxo] mimic of the ferryl intermediate in L-Arg C3 hydroxylase, VioC, reveals that HAT and -OH transfer must occur from the same side of the substrate target carbon (**A**). The distance between the reactive oxo group and the substrate target carbon (indicated by the arrows) is 3.1 Å in VioC. A structure of heme-dependent hydroxylase, P450cam, shows a similar phenomenon. A CO-bound mimic of the Fe(IV)-porphyrin radical intermediate, compound I, demands the same arrangement of HAT and -OH transfer components relative to the hydroxylation target on the camphor substrate (**B**). The distance between a mimic of the reactive group and the substrate target carbon (indicated by the arrows) is 3.1 Å in P450cam. By contrast, RS methylases use separate HAT reagents (5’-dA•) and methyl group donors (Me-Cbl), allowing for more diverse stereochemical outcomes in C-H functionalization reactions. (**C**) The structure of TokK in complex with carbapenam substrate, **1**, shows a ∼120° angle between the HAT acceptor (5’C of 5’-dAH), the substrate target carbon (C6), and the Cbl top ligand (-OH of OH-Cbl, a surrogate for Me-Cbl).

**Figure S13.**
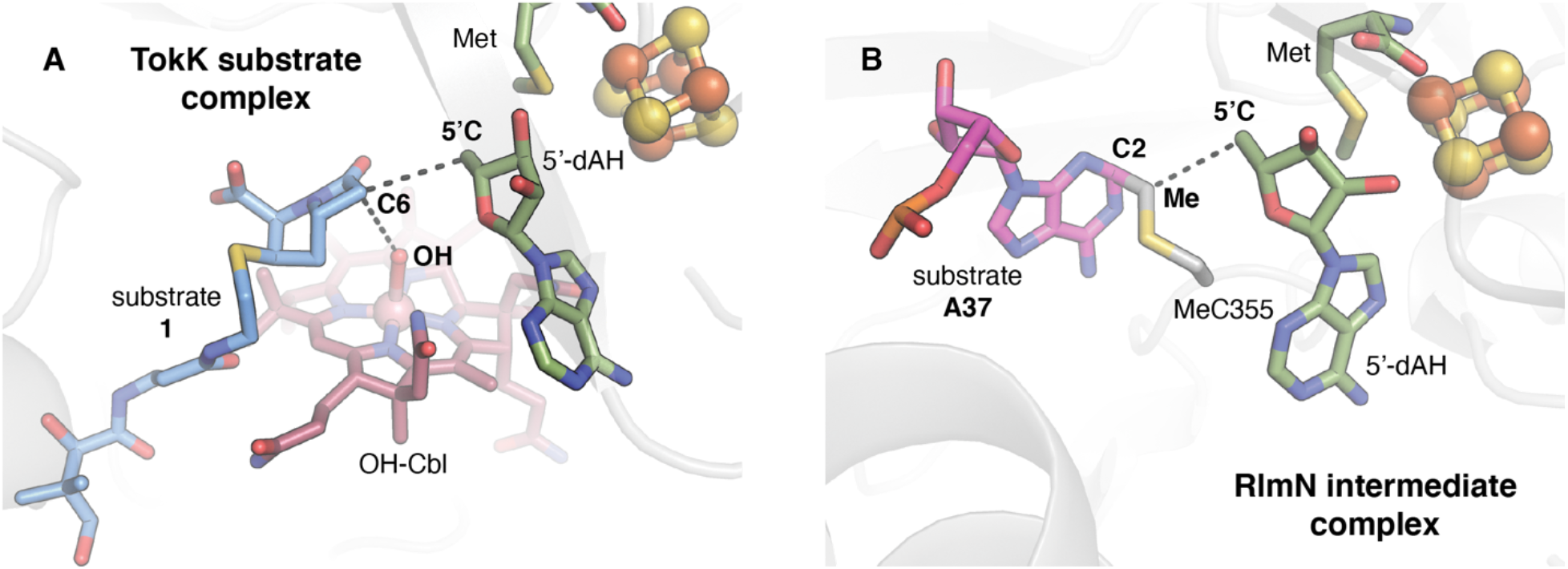
A comparison of the substrate-binding sites in two structurally characterized RS methylases. The substrate complexes of TokK (A) and RlmN (PDB ID: 5HR7) (B) are shown. RlmN is an RS methylase that uses a radical-based mechanism to methylate an *sp*^2^-hybridized carbon (C2) of an adenine base in transfer or ribosomal RNA. By contrast to TokK and other Cbl-dependent RS enzymes, RlmN uses a 5’-dA• to activate a post-translationally modified methyl-Cys residue on a C-terminal loop in the active site to modify its aromatic substrate via radical addition. Comparison of a structure of RlmN in a cross-linked methylCys-tRNA intermediate state to the TokK substrate complex shows that, despite the differences in reaction mechanism, these systems use a similar orientation of radical initiator (5’-dA•), substrate target carbon (C6, C2), and methyl donor (OH, Me). In the TokK substrate complex, the -OH ligand of the Cbl cofactor serves as a surrogate for the position of the methyl donor.

**Figure S14.**
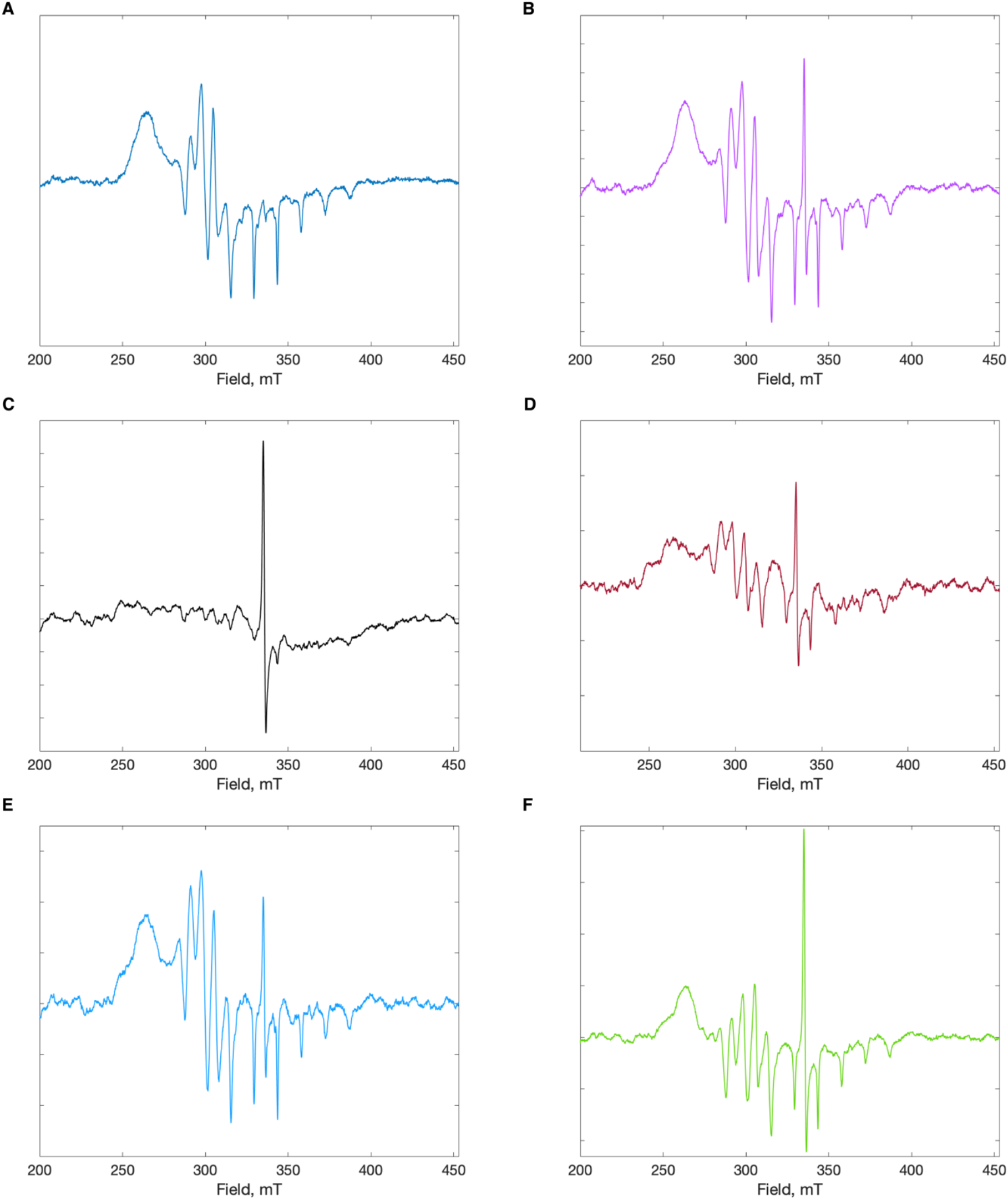
EPR analysis of the cobalamin coordination in ThnK, TokK, and TokK variants. CW-EPR spectra of ThnK (**A**), TokK (**B**), TokK W76K (**C**), TokK W76A (**D**), TokK W76F (**E**), and TokK W76H (**F**). All spectra were collected at 70 K and show the presence of 4-coordinate (or base-off, His-off) cobalamin.

**Figure S15.**
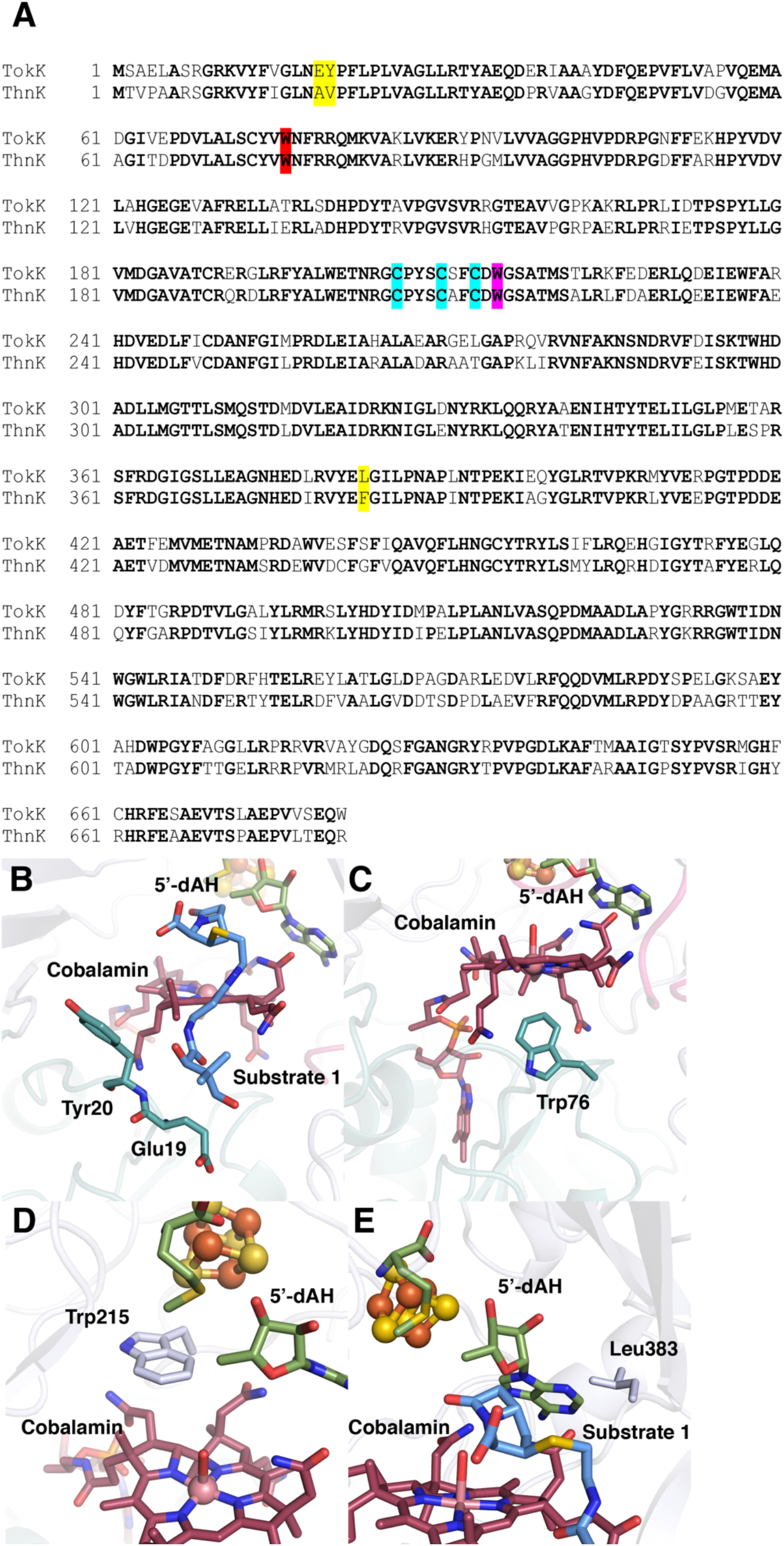
TokK and ThnK are homologous in terms of overall sequence identity but exhibit subtle differences near the carbapenem binding site. (**A**) Alignment of TokK and ThnK (uniprot accession codes A0A6B9HEI0 and F8JND9, respectively). Completely conserved residues are bolded. The conserved Trp (Trp76) beneath the cobalamin is highlighted in red. Cysteines that ligate the iron-sulfur cluster are highlighted in light blue. The Trp in the active site (Trp215) is highlighted in pink. Residues that differ between the two proteins and that were changed are highlighted in yellow. The residues that were altered are the following: Glu19 and Tyr20 (**B**), Trp76 (**C**), Trp215 (**D**), and Leu383 (**E**).

**Table S1.**
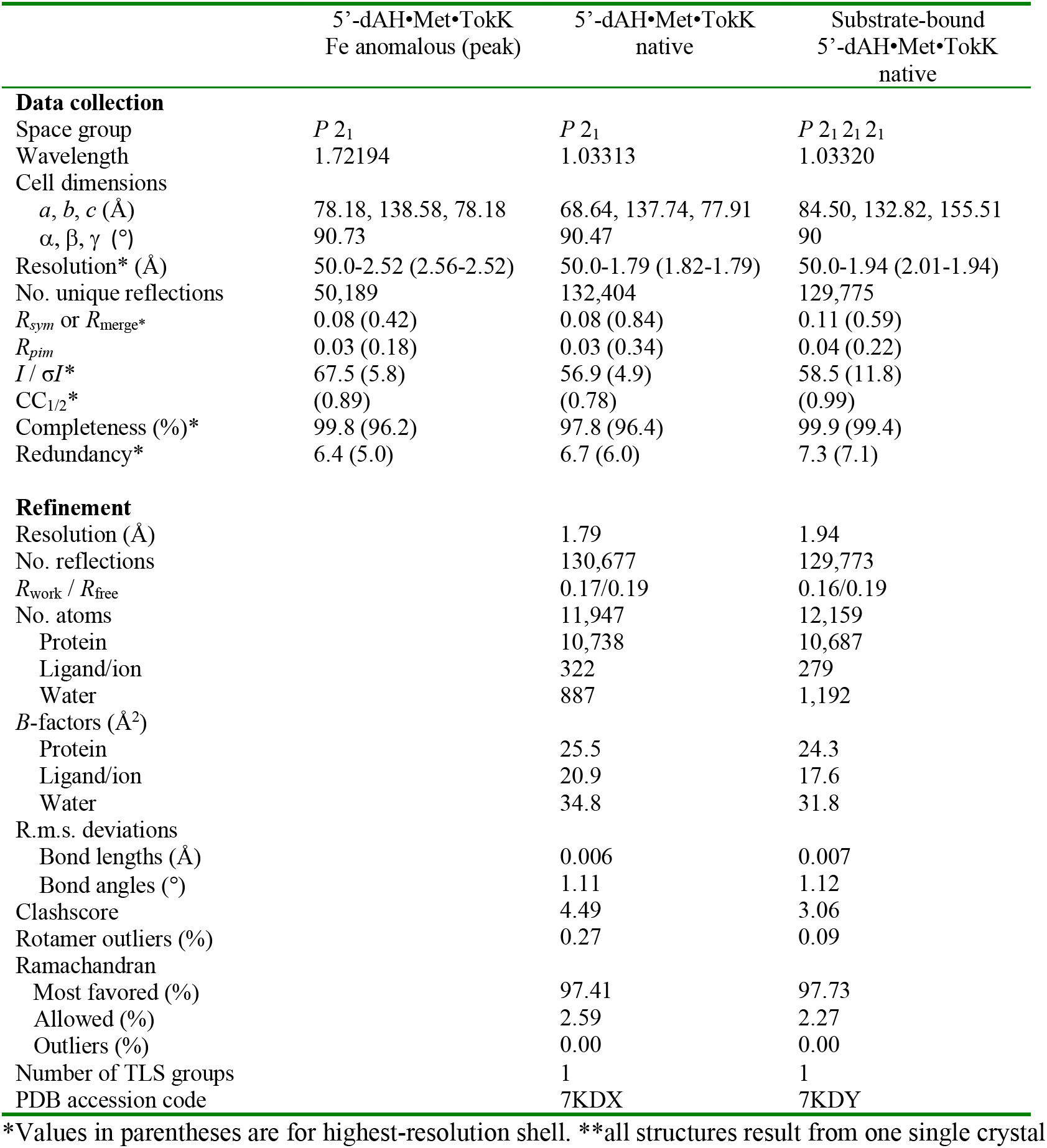
Crystallographic data table for TokK crystal structures.

**Table S2.**
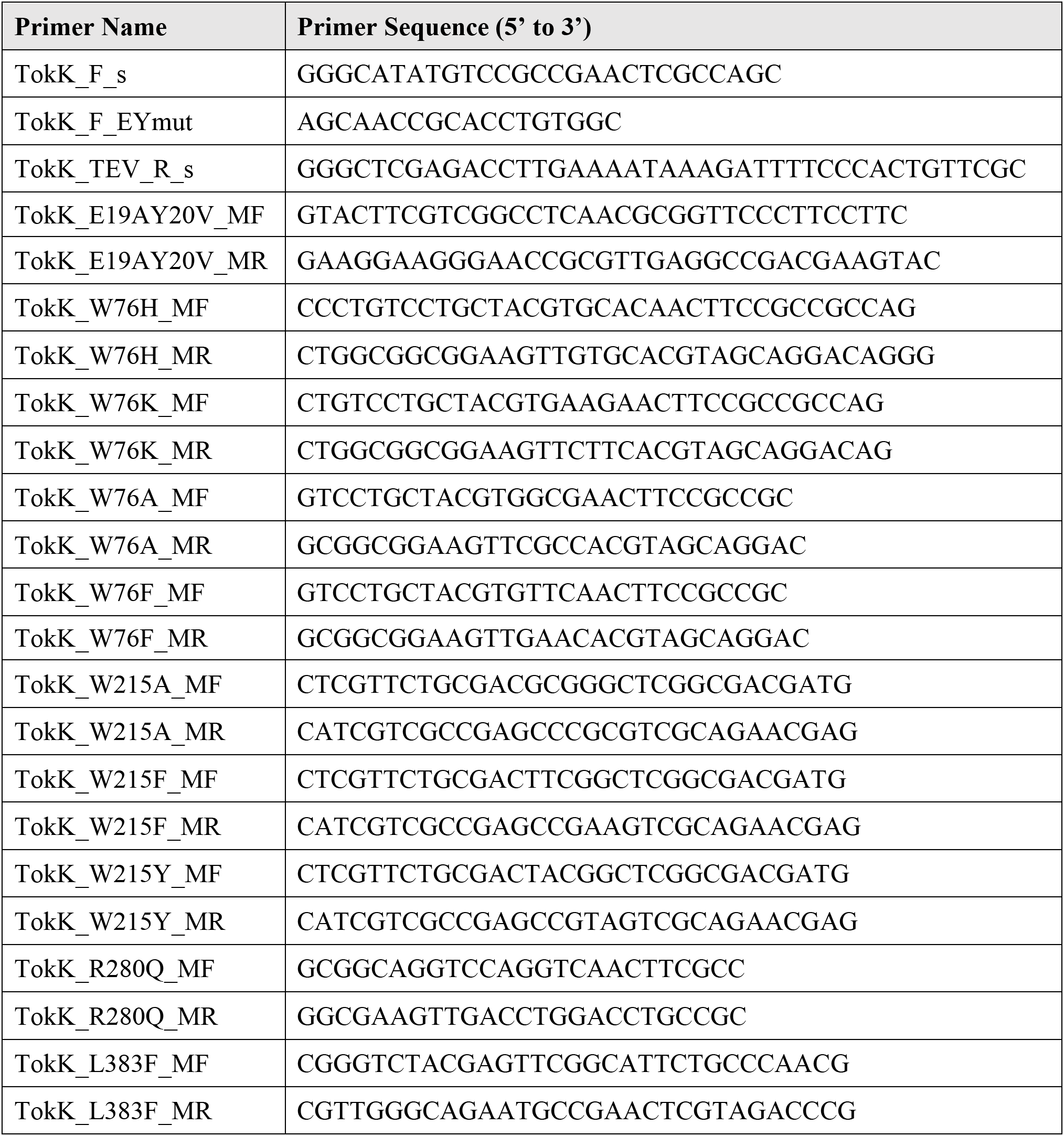
Primers used for site-directed mutagenesis of TokK

## References

1. US Food and Drug Administration, Supplemental New Drug Approval: RECARBRIO (imipenem, cilastatin, and relebactam) for injection, available at https://www.fda.gov/RegulatoryInformation/Guidances/default.htm (2020).

2. J. F. Fisher, S. Mobashery, Three decades of the class A beta-lactamase acyl-enzyme. Curr Protein Pept Sci 10, 401–407 (2009).

3. R. Li, E. P. Lloyd, K. A. Moshos, C. A. Townsend, Identification and characterization of the carbapenem MM 4550 and its gene cluster in Streptomyces argenteolus ATCC 11009. Chembiochem 15, 320–331 (2014).

4. L. E. Nunez, C. Mendez, A. F. Brana, G. Blanco, J. A. Salas, The biosynthetic gene cluster for the beta-lactam carbapenem thienamycin in Streptomyces cattleya. Chem Biol 10, 301–311 (2003).

5. S. J. Coulthurst, A. M. Barnard, G. P. Salmond, Regulation and biosynthesis of carbapenem antibiotics in bacteria. Nat Rev Microbiol 3, 295–306 (2005).

6. Y. Wang, B. Schnell, S. Baumann, R. Muller, T. P. Begley, Biosynthesis of Branched Alkoxy Groups: Iterative Methyl Group Alkylation by a Cobalamin-Dependent Radical SAM Enzyme. J Am Chem Soc 139, 1742–1745 (2017).

7. J. B. Broderick, B. R. Duffus, K. S. Duschene, E. M. Shepard, Radical S-Adenosylmethionine Enzymes. Chemical Reviews 114, 4229–4317 (2014).

8. G. L. Holliday et al., Atlas of the radical SAM superfamily: divergent evolution of function using a “plug and play” domain. Methods Enzymol 606, 1–71 (2018).

9. S. C. Wang, Cobalamin-dependent radical S-adenosyl-l-methionine enzymes in natural product biosynthesis. Nat. Prod. Rep. 35, 707–720 (2018).

10. D. V. Miller et al., in Comprehensive Natural Products III: Chemistry and Biology, H.-W. a. B. Liu, T. P., Ed. (Elsevier, 2020), vol. 5, pp. 24–69.

11. H. L. Knox et al., Structural basis for non-radical catalysis by TsrM, a radical SAM methylase. Nat Chem Biol, (2021).

12. S. Pierre et al., Thiostrepton tryptophan methyltransferase expands the chemistry of radical SAM enzymes. Nat. Chem. Biol. 8, 957–959 (2012).

13. A. J. Blaszczyk et al., Spectroscopic and Electrochemical Characterization of the Iron-Sulfur and Cobalamin Cofactors of TsrM, an Unusual Radical S-Adenosylmethionine Methylase. J Am Chem Soc 138, 3416–3426 (2016).

14. J. Bridwell-Rabb, A. Zhong, H. G. Sun, C. L. Drennan, H. W. Liu, A B12-dependent radical SAM enzyme involved in oxetanocin A biosynthesis. Nature 544, 322–326 (2017).

15. A. J. Blaszczyk, B. Wang, A. Silakov, J. V. Ho, S. J. Booker, Efficient methylation of C2 in L-tryptophan by the cobalamin-dependent radical S-adenosylmethionine methylase TsrM requires an unmodified N1 amine. J. Biol. Chem. 292, 15456–15467 (2017).

16. W.-c. Chang et al., Mechanism of the C5 stereoinversion reaction in the biosynthesis of carbapenem antibiotics. Science 343, 1140–1144 (2014).

17. E. K. Sinner, M. S. Lichstrahl, R. Li, D. R. Marous, C. A. Townsend, Methylations in complex carbapenem biosynthesis are catalyzed by a single cobalamin-dependent radical S-adenosylmethionine enzyme. Chem Commun (Camb) 55, 14934–14937 (2019).

18. G. A. Albers-Schoenberg, Byron H.; Hensens, Otto D.; Hirshfield, Jordan; Hoogsteen, Karst; Kaczka, Edward A.; Rhodes, Robert E.; Kahan, Jean S.; Kahan, Frederick M.; Ratcliff, Ronald W.; Walton, Edward; Ruswinkle, Linda J.; Morin, Robert B.; and Christensen, Burton G, Structure and absolute configuration of thienamycin. J Am Chem Soc 100, 6491–6499 (1978).

19. A. J. Mitchell et al., Visualizing the Reaction Cycle in an Iron(II)- and 2-(Oxo)-glutarate-Dependent Hydroxylase. J Am Chem Soc 139, 13830–13836 (2017).

20. T. L. Poulos, B. C. Finzel, A. J. Howard, High-resolution crystal structure of cytochrome P450cam. J Mol Biol 195, 687–700 (1987).

21. E. L. Schwalm, T. L. Grove, S. J. Booker, A. K. Boal, Crystallographic capture of a radical S-adenosylmethionine enzyme in the act of modifying tRNA. Science 352, 309–312 (2016).

22. C. L. Drennan, S. Huang, J. T. Drummond, R. G. Matthews, M. L. Ludwig, How a Protein Binds B-12 -a 3.0-Angstrom X-Ray Structure of B-12-Binding Domains of Methionine Synthase. Science 266, 1669–1674 (1994).

23. B. Kräutler, Thermodynamic *trans*-effects of the nucleotide base in the B12 coenzymes. Helv. Chim. ACTA 70, 1268–1278 (1987).

24. B. Krautler, Chemistry of methylcorrinoids related to their roles in bacterial C1 metabolism. FEMS Microbiol Rev 7, 349–354 (1990).

25. P. A. Milton, T. L. Brown, The kinetics of benzimidazole dissociation in methylcobalamin. J Am Chem Soc 99, 1390–1396 (1977).

26. D. R. Marous et al., Consecutive radical S-adenosylmethionine methylations form the ethyl side chain in thienamycin biosynthesis. Proc. Natl. Acad. Sci. 112, 10354–10358 (2015).

27. N. D. Lanz et al., Enhanced Solubilization of Class B Radical S-Adenosylmethionine Methylases by Improved Cobalamin Uptake in Escherichia coli. Biochemistry 57, 1475–1490 (2018).

28. N. D. Lanz et al., RImN and AtsB as Models for the Overproduction and Characterization Radical SAM Proteins. *Natural Product Biosynthesis by Microorganisms and Plants*, Pt B 516, 125–152 (2012).

29. A. J. Blaszczyk, R. X. Wang, S. J. Booker, TsrM as a Model for Purifying and Characterizing Cobalamin-Dependent Radical S-Adenosylmethionine Methylases. Methods Enzymol. 595, 303–329 (2017).

30. P. D. Adams et al., PHENIX: a comprehensive Python-based system for macromolecular structure solution. Acta Crystallographica Section D-Biological Crystallography 66, 213–221 (2010).

31. G. Bunkoczi, et al., Phaser.MRage: automated molecular replacement. Acta Crystallographica Section D-Biological Crystallography 69, 2276–2286 (2013).

32. W. Minor, M. Cymborowski, Z. Otwinowski, M. Chruszcz, HKL-3000: the integration of data reduction and structure solution -from diffraction images to an initial model in minutes. Acta Crystallographica Section D-Biological Crystallography 62, 859–866 (2006).

33. Z. Otwinowski, W. Minor, Processing of X-ray diffraction data collected in oscillation mode. Macromolecular Crystallography, Pt A 276, 307–326 (1997).

34. P. Emsley, B. Lohkamp, W. G. Scott, K. Cowtan, Features and development of Coot. Acta Crystallographica Section D-Biological Crystallography 66, 486–501 (2010).

35. V. B. Chen et al., MolProbity: all-atom structure validation for macromolecular crystallography. Acta Crystallographica Section D-Structural Biology 66, 12–21 (2010).

36. The PyMOL Molecular Graphics Systems, Version 2.0 Schrödinger, LLC.

37. B. K. Ho, F. Gruswitz, HOLLOW: generating accurate representations of channel and interior surfaces in molecular structures. BMC Struct Biol 8, 49 (2008).

38. A. C. Wallace, R. A. Laskowski, J. M. Thornton, LIGPLOT: a program to generate schematic diagrams of protein-ligand interactions. Protein Eng 8, 127–134 (1995).

39. A. W. Schuttelkopf, D. M. van Aalten, PRODRG: a tool for high-throughput crystallography of protein-ligand complexes. Acta Crystallogr D Biol Crystallogr 60, 1355–1363 (2004).

40. E. Akiva et al., The Structure-Function Linkage Database. Nucleic Acids Research 42, D521–D530 (2014).

41. J. A. Gerlt, Genomic Enzymology: Web Tools for Leveraging Protein Family Sequence-Function Space and Genome Context to Discover Novel Functions. Biochemistry 56, 4293–4308 (2017).

42. H. J. Kim et al., GenK-catalyzed C-6’ methylation in the biosynthesis of gentamicin: isolation and characterization of a cobalamin-dependent radical SAM enzyme. J. Am. Chem. Soc. 135, 8093–8096 (2013).

43. C. Huang et al., Delineating the biosynthesis of gentamicin x2, the common precursor of the gentamicin C antibiotic complex. Chem Biol 22, 251–261 (2015).

44. W. J. Werner et al., In Vitro Phosphinate Methylation by PhpK from Kitasatospora phosalacinea. Biochemistry 50, 8986–8988 (2011).

45. B. Wang et al., Stereochemical and Mechanistic Investigation of the Reaction Catalyzed by Fom3 from Streptomyces fradiae, a Cobalamin-Dependent Radical S-Adenosylmethionine Methylase. Biochemistry 57, 4972–4984 (2018).

46. A. Parent et al., The B12-Radical SAM Enzyme PoyC Catalyzes Valine Cβ-Methylation during Polytheonamide Biosynthesis. J. Am. Chem. Soc. 138, 15515–15518 (2016).

47. K. Watanabe et al., Escherichia coli allows efficient modular incorporation of newly isolated quinomycin biosynthetic enzyme into echinomycin biosynthetic pathway for rational design and synthesis of potent antibiotic unnatural natural product. J Am Chem Soc 131, 9347–9353 (2009).

48. J. Y. Suzuki, D. W. Bollivar, C. E. Bauer, Genetic analysis of chlorophyll biosynthesis. Annu Rev Genet 31, 61–89 (1997).

49. L. Westrich, L. Heide, S. M. Li, CloN6, a novel methyltransferase catalysing the methylation of the pyrrole-2-carboxyl moiety of clorobiocin. ChemBioChem 4, 768–773 (2003).

50. M. I. Radle, D. V. Miller, T. N. Laremore, S. J. Booker, Methanogenesis marker protein 10 (Mmp10) from Methanosarcina acetivorans is a radical S-adenosylmethionine methylase that unexpectedly requires cobalamin. Journal of Biological Chemistry 294, 11712–11725 (2019).

51. J. A. Gerlt et al., Enzyme Function Initiative-Enzyme Similarity Tool (EFI-EST): A web tool for generating protein sequence similarity networks. Biochim Biophys Acta 1854, 1019–1037 (2015).

52. R. Zallot, N. Oberg, J. A. Gerlt, The EFI Web Resource for Genomic Enzymology Tools: Leveraging Protein, Genome, and Metagenome Databases to Discover Novel Enzymes and Metabolic Pathways. Biochemistry 58, 4169–4182 (2019).

53. R. Zallot, N. O. Oberg, J. A. Gerlt, ’Democratized’ genomic enzymology web tools for functional assignment. Curr Opin Chem Biol 47, 77–85 (2018).

54. P. Shannon et al., Cytoscape: a software environment for integrated models of biomolecular interaction networks. Genome Res 13, 2498–2504 (2003).

55. S. Hoops et al., COPASI--a COmplex PAthway SImulator. Bioinformatics 22, 3067–3074 (2006).

56. J. Schaff, C. C. Fink, B. Slepchenko, J. H. Carson, L. M. Loew, A general computational framework for modeling cellular structure and function. Biophys J 73, 1135–1146 (1997).

